# The role of Platelet-derived growth factor (PDGF) in uremic vascular calcification

**DOI:** 10.1101/2025.07.09.664016

**Authors:** Burcu Yesilyurt Öztürk, Li Zhang, Saskia von Stillfried, Barbara Mara Klinkhammer, Julia Möllmann, Patrick Droste, Eva Miriam Buhl, Mathias Hohl, Heidi Noels, Vera Jankowski, Joachim Jankowski, Claudia Goettsch, Nikolaus Marx, Lorin E. Olson, Jürgen Floege, Dickson W.L. Wong, Peter Boor

## Abstract

Vascular calcification is common in chronic kidney disease (CKD), contributing to increased cardiovascular morbidity and mortality. One of the proposed mechanisms of driving vascular calcification is a phenotypic switch of vascular smooth muscle cells (VSMCs). The platelet-derived growth factors (PDGFs) and their receptors (PDGFRs), particularly PDGFR-β, were shown to modulate the VSMC phenotype. However, their role in uremic vascular calcification remained unclear.

We adapted an *ex vivo* calcification model using murine aortas to simulate uremic conditions. Compared to control conditions, incubation with hemodialysate from CKD patients or using aortas from CKD animals both resulted in significantly increased PDGFR-β phosphorylation and vascular calcification. Inhibition of PDGF signaling using soluble PDGFR-β or the small molecule tyrosine kinase inhibitor imatinib significantly reduced uremic calcification and enhanced vascular elasticity. Next, we generated transgenic mice with a VSMC-specific, inducible expression of constitutively active PDGFR-β. The aortas of these mice exhibited significantly increased vascular calcification *ex vivo*, which was further aggravated by uremic conditions. We established an *in vivo* model of accelerated vascular calcification and CKD in the transgenic mice, showing significantly aggravated vascular calcification and phenotypic switching of VSMCs compared to non-transgenic littermates. Finally, increased expression of phosphorylated PDGFR-β and a VSMC phenotypic switching were detected in human arteries from patients with CKD compared to those without CKD.

In conclusion, PDGFR-β contributes to CKD-associated vascular calcification, representing a potential novel therapeutic target.

## Introduction

In comparison to the general population, cardiovascular morbidity and mortality are significantly elevated in chronic kidney disease (CKD) patients, with risk increasing progressively across CKD stages [1]. Vascular calcification is a major complication in advanced CKD, affecting up to 50% of patients with CKD stage 4–5 [2], and 80% of those with end-stage kidney disease [3]. Medial calcification, characterized by excessive calcium and phosphate deposition within the media of arteries, results in arterial stiffness and diminished vascular elasticity [4, 5]. This calcification increases systolic blood pressure, promotes left ventricular hypertrophy, and is associated with heart failure [6–8]. Vascular calcification is an active, cell-regulated process driven by vascular smooth muscle cells (VSMCs) [9]. Under physiological conditions, VSMCs maintain a contractile phenotype, but CKD-associated accumulation of phosphate and other uremic toxins promotes their phenotypic transition to a synthetic state and subsequent matrix mineralization [10]. This phenotypic switch involves the downregulation of contractile markers such as smooth muscle (SM) 22α, alongside the upregulation of osteochondrogenic markers such as osteopontin (OPN) [11, 12].

Uremic conditions were shown to induce VSMC transition from a contractile to a synthetic state, both *in vitro* and *in vivo* [13]. *In vitro*, uremic serum enhanced VSMC mineralization partially via increased OPN [14, 15].

Platelet-derived growth factor receptor β (PDGFR-β) belongs to the PDGF protein family, consisting of five dimeric ligands and two tyrosine kinase receptor chains. The PDGF protein family orchestrates PDGF signaling in vascular development and pathology. PDGFR-β, typically expressed at low activity in VSMCs, becomes upregulated during disease states [16]. PDGFR-β is present in atherosclerotic lesions and contributes to vasculopathy through VSMC proliferation, migration, induction of inflammation, and/or oxidative stress [17].

A transgenic mouse model with activated PDGFR-β in VSMCs accelerated atherosclerosis, tunica media thickening, and VSMC phenotypic switch by suppressing the contractile genes [18]. *In vitro* studies suggested that VSMCs exposed to a characteristic uremic toxin (indoxyl sulfate), exhibited increased PDGFR-β phosphorylation and subsequent VSMC proliferation and migration, contributing to the development of atherosclerosis [19]. While PDGFR-β activation in VSMCs is known to contribute to vasculopathy by modulating VSMC phenotype, its specific role in CKD-associated vascular calcification remains largely unexplored.

In this study, we investigated the role of PDGFR-β in uremic vascular calcification using novel *ex vivo* and *in vivo* CKD-associated vascular calcification models and a transgenic mouse line. We addressed the translational relevance in a cohort of human artery samples. **Our results indicate PDGFR-β activation in VSMCs as a novel mediator of vascular calcification in CKD.**

## Methods

### Animals for *ex vivo* aorta culture and *in vivo* CKD models

All animal experiments were approved by the Institute of Laboratory Animal Science at RWTH Aachen University Hospital and the State Office for Consumer Protection and Food (*Landesamt für Verbraucherschutz und Ernährung NRW*) in accordance with Section 11 of the German Animal Welfare Act (TSchG). Procedures complied with the German Animal Welfare Ordinance on Experimental Animals, EU Directive 2010/63, and the recommendations of FELASA and GV-SOLAS. All experiments followed the ARRIVE (Animal Research: Reporting of *In Vivo* Experiments) guidelines and were conducted in accordance with the guidelines for the Care and Use of Laboratory Animals.

Mice were housed in an ISO 9001:2015-certified facility under controlled conditions (20– 24 °C, 55% ± 10% humidity, 12-hour light/dark cycle) with *ad libitum* access to food and water. Upon sacrifice, mice were anesthetized using either isoflurane (overdosing) or intraperitoneal injection of ketamine (100 mg/ body weight kg) and xylazine (10 mg/ body weight kg). Aortas were harvested from the carotid arteries to the femoral bifurcation, and surrounding connective tissues were cleaned before *ex vivo* culturing.

- Age, sex, background, and aortic region comparison for *ex vivo* experiments Wild-type (WT) mice of both sexes, aged 10 up to 80 weeks, and from various genetic backgrounds (C57BL/6J, SV129, mixed C57BL/6–SV129, and FVB) were used. Aortas were cultured *ex vivo* in either growth or calcification medium for up to 10 days. Incubation durations ranged from 3 to 10 days, and distinct aortic regions (arch, thoracic, suprarenal, and infrarenal) were analyzed separately.
- Aortas from CKD mice for *ex vivo* experiments 13 C57BL/6NRj male mice (Janvier Labs, France) were randomly assigned to two groups (N = 6 standard diet, N = 7 adenine diet) and fed either a standard diet or an adenine-enriched diet containing 0.1% or 0.2% alternating adenine (ssniff Spezialdiäten GmbH, Soest, Germany) for 8 weeks to induce 2,8-dihydroxyadenine nephropathy, as previously described [20]. Aortas from CKD mice were harvested and cultured *ex vivo* in either growth or calcification medium for 7 days, respectively (see Aorta removal and *ex vivo* culture).
- Myh11-Cre PDGFR-βV536A mutant mice (VRbJ) for *ex vivo* and *in vivo* experiments Myh11-CreER^T2^ mice (The Jackson Laboratory JAX® No. 019079) were crossed with PDGFR-βV536A mutant mice [18] (Myh11Cre+::Pdgfrb^+/J^; VRbJ) to generate a murine model with tamoxifen-inducible PDGFR-β activation specifically in VSMCs, with expression of the constitutively active PDGFR-β driven by the endogenous PDGFR-β promoter. Mice were backcrossed to SV129X1 background. Wild-type (Myh11Cre-::Pdgfrb^+/+^) and Cre+ (Myh11Cre+::Pdgfrb^+/+^) littermates were used as the control/non-mutant group (WT) from which aortas were compared to VRbJ mice aortas.
- PDGFR-β reporter mouse model for *in vivo* experiments PDGFR-β-eGFP (FVB/N-Tg) reporter mice were obtained from GENSAT (https://www.mmrrc.org/catalog/sds.php?mmrrc_id=31796). A total of 5 mice were fed a standard diet and sacrificed at baseline (week 0) and 10 mice were subjected to an *in vivo* CKD-calcification model (Suppl. Figure 3A). Mice aged 8–14 weeks were included in the studies.

### Aorta removal and *ex vivo* culture

Aortas were incubated in either a growth or an adapted calcification medium for different durations (3, 5, 7, and 10 days). The growth medium consisted of Dulbecco’s Modified Eagle Medium (DMEM) supplemented with 10% fetal bovine serum (FBS), 1% penicillin/streptomycin, and gentamicin (all from Thermo Fisher Scientific, Waltham, MA). The calcification medium contained additional supplements, including 10 mmol/L β-glycerophosphate, 8 mmol/L CaCl₂, 10 mmol/L sodium pyruvate, 50 µg/mL L-ascorbic acid, and 100 nmol/L dexamethasone. The culture medium was refreshed every other day. Recombinant mouse PDGFR-β Fc chimera (R&D Systems, Cat# 1042-PR) and Imatinib (Sigma-Aldrich, Cat# SML1027, St. Louis, MO) were administered at final concentrations of 0.5 µg/mL and 1 or 2 µM, respectively.

For the aortas harvested from VRbJ (Myh11Cre+::Pdgfrb^+/J^) and WT (Myh11Cre-::Pdgfrb^+/+^ and Myh11Cre+::Pdgfrb^+/+^) mice, fresh 4’OH-tamoxifen (1 µM; Sigma-Aldrich, Cat# 176141, St. Louis, MO) was applied 4 times over the 7-day culture period (on days 1, 2, 3, and 5) to activate the Cre recombinase.

Our *ex vivo* CKD model was established by the stimulation with human CKD hemodialysate. Pooled hemodialysate fractions were kindly provided by the Institute for Molecular Cardiovascular Research (IMCAR), RWTH Aachen University, Aachen, Germany [21]. Ethical approval was obtained from local authorities (EK/196/18). Briefly, hemodialysate collected from CKD patients was fractionated and concentrated using reverse-phase chromatography (LiChroprep® RP-18 (40–63 µm); Merck) and eluted based on the substance solubility present in the hemodialysate, followed by lyophilization and reconstitution in 5 mL ddH_2_O. Hemodialysate fractions were pooled to represent the full spectrum of circulating uremic toxins present in CKD patients and added to cell culture medium in 1:250 dilution.

### CKD-calcification *in vivo* model

To induce CKD-associated vascular calcification, mice received a tail vein injection of adeno-associated virus encoding gain-of-function PCSK9 (AAV8-D377Y-mPCSK9, 10^11^ virus particles, Vector Biolabs, PA, USA), promoting extracellular calcium deposition [22]. Following injection, mice were fed a mixed diet for 12 weeks, consisting of 0.2% adenine, 1.8% phosphate, and Western diet (21% high-fat, 0.21% high-cholesterol, ssniff Spezialdiäten GmbH, Soest, Germany) (Figure 5A, suppl. Figure 3A).

PDGFR-β reporter mice (FVB/N-Tg Pdgfrb-eGFP, N = 10, 4 male and 6 female) were subjected to the CKD calcification model compared to the control mice (N = 5, male) fed a standard diet and sacrificed at baseline (week 0) for comparison.

For VRbJ and WT littermates, tamoxifen (30 mg/mL, Merck Sigma-Aldrich, Cat# 10540-29-1, St. Louis, MO) was first administered intraperitoneally 3 times a week (day −14, −12, −10) in advance of the *in vivo* CKD-calcification model. Tamoxifen was dissolved in 97% corn oil (sterile filtered) and 3% ethanol. AAV8-D377Y-mPCSK9 was injected into both non-mutant (WT: Myh11Cre-::Pdgfrb^+/+^ and Myh11Cre+::Pdgfrb^+/+^; N = 12, 7 male and 5 female) and mutant (VRbJ: Myh11Cre+::Pdgfrb^+/J^; N = 8, male) mice at day 0 (Figure 5A). As the Cre gene is Y chromosome-linked, all Cre+ mice were male. Mice were maintained on the same diet for 12 weeks. Urine was collected in metabolic cages for 12-16 hours overnight at the end point for further analysis. OsteoSense™ 680 EX was administered via tail vein injection 48 hours before sacrifice for visualizing tissue calcification.

After deep anesthesia upon the sacrifice, retroorbital blood sampling was performed, followed by a whole-body perfusion with 0.9% saline via the heart apex as described [23]. Kidneys and heart connected with aorta were harvested *en bloque* for OsteoSense imaging with Odyssey® Sa Imager model 9260 (LI-COR Biosciences, USA) and immediately processed for further histological and molecular analysis as described below. Additional organs, including the liver, were collected and processed similarly. Metabolites (e.g., creatinine and urea) were analyzed in the serum and urine by Vitros 350 Chemistry Analyzer (Ortho Clinical Diagnostics, Unterschleißheim, Germany).

### Blood pressure and electrocardiography (ECG) measurements

Non-invasive blood pressure was measured using the CODA Volume Pressure Recording (VPR) system (Kent Scientific Corporation, USA), which utilizes an occlusion tail-cuff sensor and mice were stabilized on a warming platform (32–35°C) during the measurement. Heart rate and electrocardiographic signals were recorded in conscious, unrestrained mice using the ECGenie system (Mouse Specifics, Inc., USA). Mice were acclimated for around five minutes before measurements to minimize stress-related variability. Data acquisition was performed using LabChart software (ADInstruments, New Zealand), and ECG signals were analyzed using the eMouse program (iWorx Systems Inc., USA).

### Calcium measurement

The calcium content of aortic tissue was quantified using the Randox Calcium Assay Kit (Randox Laboratories, CA590). In short, aortas were incubated in 0.6 N HCl for 24 hours at 4°C, and dissolved Ca^2+^ ions were quantified following the manufacturer’s protocol. Absorbance was measured at 570 nm using a microplate reader (Tecan, Switzerland).

Following calcium measurement, the same aortic tissues were solubilized in 0.1 N NaOH/0.1% sodium dodecyl sulfate (SDS) (Carl Roth). Total protein content was determined using the Bicinchoninic Acid (BCA) Assay Kit (Interchim, UP40840A). Calcium levels were normalized to total protein, and results were expressed as a fold change relative to control.

### RNA isolation and Real-Time quantitative PCR (RT-qPCR)

Total RNA and protein were extracted from aortic tissues using the NucleoSpin® TriPrep Isolation Kit (Macherey-Nagel, Germany) following the manufacturer’s instructions. Complementary DNA (cDNA) was synthesized from the RNA with M-MLV (Moloney-murine leukemia virus-reverse transcriptase) enzyme (Invitrogen), followed by RT-qPCR performed with a CFX Opus 96 system (Bio-Rad, Hercules, CA). cDNA samples were mixed with iTaq™ Universal SYBR® Green Supermix (Bio-Rad) and the primer pairs listed in Table 1. Data was normalized to *Gapdh*.

**Table 1:**
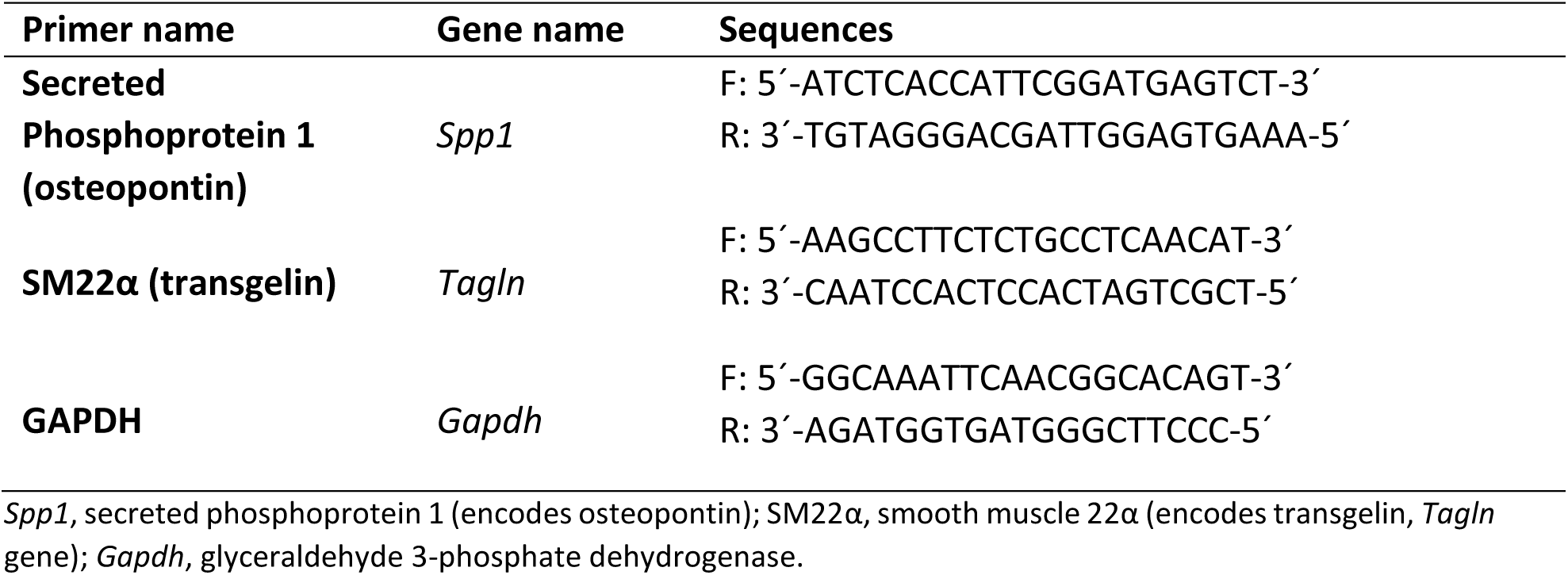
Primers used in qPCR.

### Protein isolation and Western Blot

Protein isolation was followed by the RNA isolation with NucleoSpin® TriPrep Isolation Kit, according to the manufacturer’s manual. Protein concentrations were measured by a quantification assay kit (Macherey-Nagel, Germany) and equal amounts of protein from each sample were resolved in NuPAGE 4-12% Bis-Tris gels and transferred onto 0.2 or 0.45 µm nitrocellulose membranes (GE Healthcare) with the Mini Bolt™ semi-dry transfer system (Invitrogen). Membranes were then blocked with 3% Bovine Serum Albumin (BSA, Serva, Germany) for 1 hour. Primary antibodies were incubated overnight at 4°C, followed by 1-hour incubation with secondary antibodies (Table 2-3). Protein bands were visualized by an Odyssey® Sa Imager model 9260 (LI-COR Biosciences, USA) and iBright™ FL1500 Imaging System (Invitrogen) and analyzed by iBright Analysis Software (Invitrogen).

**Table 2:** Primary Antibodies for staining (IHC/IF) and protein analysis (WB)

| Name | Host species | Supplier (Catalog-No.) | Dilution (IHC/IF) | Dilution (WB) |
| --- | --- | --- | --- | --- |
| **p-PDGFR-β Tyr 1009** | mouse (monoclonal) | Santa Cruz Biotechnology, Inc. (sc-373805) | 1:100 |  |
| **p-PDGFR-β Tyr 1009** | rabbit (monoclonal) | Cell Signaling Technology (3124) |  | 1:1000 |
| **Cre Recombinase** | rabbit (monoclonal) | Cell Signaling Technology (15036) | 1:100 | 1:1000 |
| **Osteopontin** | goat (polyclonal) | R&D Systems (AF808) | 1:200 | 1:1000 |
| **Actin** | rabbit (polyclonal) | Sigma-Aldrich (A2066) |  | 1:2000 |
| **LDL Receptor** | rabbit (monoclonal) | Abcam (ab52818 [EP1553Y]) |  | 1:1000 |
| **α-SMA** | mouse (monoclonal) | Dako (M0851) | 1:100 |  |

p-PDGFR-β Tyr 1009, phosphorylated platelet-derived growth factor receptor $\beta$ at tyrosine residue 1009; LDL receptor, low density lipoprotein receptor; α-SMA, alpha-smooth muscle actin.

**Table 3:** Secondary Antibodies for staining (IHC/IF) and protein analysis (WB)

| Name | Host species | Supplier (Catalog-No.) | Dilution (IHC/IF) | Dilution (WB) |
| --- | --- | --- | --- | --- |
| **anti-rabbit** | goat | Vector (BA-1000) | 1:300 |  |
| **anti-mouse** | horse | Vector (BA-2001) | 1:300 |  |
| **anti-goat** | donkey | Invitrogen (A16005) | 1:300 |  |
| **anti-mouse Alexa Fluor Plus 488** | goat | Invitrogen (A48286) | 1:100 |  |
| **anti-rabbit Alexa Fluor Plus 488, 555, 647** | goat | Invitrogen (A48285) |  | 1:10000 |
| **anti- goat Alexa Fluor Plus 488, 555, 647** | donkey | Invitrogen (A32816) |  | 1:10000 |

### Histology

#### Tissue Collection, Processing, and Analysis

Ethical approval for analysis of archived human formalin-fixed, paraffin-embedded (FFPE) samples was obtained from local authorities (EK/042/17, EK/294/22). In the laboratory information system at the Institute of Pathology at RWTH Aachen, N=5 patients with chronic kidney disease, stage 5 (ICD-10 N18.5), with calcified arteries in amputation specimens and N=5 FFPE samples from age- and sex matched non-CKD patients with calcified arteries in amputation specimens were identified. Next, N=5 age- and sex matched patients with CKD, stage 5, with non-calcified arteries in amputation specimens and N=6 FFPE samples from age- and sex matched non-CKD patients with non-calcified arteries in amputation specimens were identified, respectively (Table 4). The selected FFPE tissue from amputated extremities was dearchived from the diagnostic archives at the Institute of Pathology at RWTH Aachen University Hospital. FFPE blocks were decalcified *en bloque* in EDTA for seven days prior to cutting. 4 µm sections were prepared for staining and histological analysis.

**Table 4:** Characteristics of CKD and non-CKD patients with calcified and non-calcified arteries.

| Variable | CKD-calcified (n=5) | CKD-non-calcified (n=5) | Non-CKD-calcified (n=5) | Non-CKD-non-calcified (n=6) |
| --- | --- | --- | --- | --- |
| **Sex, male** | 4/5 (80%) | 3/5 (60%) | 4/5 (80%) | 4/6 (66%) |
| **Age, years** | 64.8 ± 7.3 | 67.0 ± 13.7 | 65.6 ± 7.4 | 37.3 ± 6.6 |
| **BMI, kg/m<sup>2</sup>** | 28.9 ± 6.6 | 25.7 ± 10.6 | 27.5 ± 7.5 | 24.9 ± 2.5 |
| **Artery involvement** |  |  |  |  |
| - Popliteal artery | 1/5 (20%) | 0/5 (0%) | 1/5 (20%) | 2/6 (33%) |
| - Anterior and posterior tibial artery | 3/5 (60%) | 3/5 (60%) | 4/5 (80%) | 1/6 (16%) |
| - Other arteries (medial malleolar, ulnar/radial, or not specified) | 1/5 (20%) | 2/5 (40%) | 0/5 (0%) | 3/6 (50%) |
| **Indications for surgery** |  |  |  |  |
| - Peripheral arterial disease | 3/5 (60%) | 1/5 (20%) | 2/5 (40%) | 0/6 (0%) |
| - Gangrene | 1/5 (20%) | 1/5 (20%) | 1/5 (20%) | 1/6 (16%) |
| - Infection/necrosis | 1/5 (20%) | 1/5 (20%) | 0/5 (0%) | 2/6 (33%) |
| - Other (diabetic foot, osteomyelitis, ischemia, ulcers, tumors, or not specified) | 0/5 (0%) | 2/5 (40%) | 2/5 (40%) | 3/6 (50%) |

Baseline characteristics of patients divided into four groups based on chronic kidney disease (CKD) status and vascular calcification status: CKD-calcified (n=5), CKD-non-calcified (n=5), non-CKD-calcified (n=5), and non-CKD-non-calcified (n=6). The table summarizes demographic variables (sex, age, BMI), artery involvement, and indications for surgery. Data are presented as continuous variables as mean ± SD, while categorical variables are expressed as n/N (%).

Similarly, *ex vivo* murine aortas were fixed in 4% paraformaldehyde (PFA, Morphisto, Germany) and *in vivo* murine aortas were fixed with 4% PFA or methyl Carnoy’s solution, followed by tissue processing and paraffin embedding. 1 µm sections were used for histological stainings.

To analyze the stained sections, all slides were either scanned with a whole-slide scanner (AT2, Leica Biosystems, Wetzlar, Germany) or imaged by the Zeiss Axio Imager 2 fluorescence microscope (Carl Zeiss, Germany). Image analysis (positively stained pixels in the region of interest, i.e., tunica media) was performed using ImageJ or a custom macro in ImageScope (Carl Zeiss, Germany). The percentage of human artery calcification was calculated as the ratio of calcified area to the total arterial area.

### Histological staining, immunohistochemistry (IHC) and immunofluorescence (IF)

Kidney tissues were stained with Periodic Acid–Schiff (PAS) reagent (Merck, Germany) and Acid Fuchsin Orange G (AFOG, Sigma-Aldrich, St. Louis, MO) as described before [24] for evaluating kidney pathologies and fibrosis, respectively. Human artery samples were stained with hematoxylin and eosin (H&E) solution (Sigma-Aldrich, St. Louis, MO) for pathological features as described [25]. Von Kossa [26, 27], Alizarin red and OsteoSense stainings were performed for visualizing vascular calcification. Alizarin Red and von Kossa are well-established histological techniques for detecting macrocalcifications, primarily reflecting calcium and phosphate deposits, respectively [28]. In contrast, OsteoSense, a near-infrared fluorescent probe, binds specifically to hydroxyapatite with high sensitivity, enabling the detection of microcalcifications across *in vitro*, *ex vivo*, and *in vivo* models [28–30].

For aortic sections, 2% Alizarin Red S (Sigma-Aldrich, A5533, St. Louis, MO) was applied for 4 minutes, followed by a washing step and dehydration sequentially in acetone, acetone-xylene (1:1), and xylene before mounting. For whole aortas, tissues were incubated in 0.0016% Alizarin Red S dissolved in 0.5% KOH (Sigma-Aldrich, St. Louis, MO) at room temperature for 24 hours, followed by washing with 0.05% KOH for an additional 24 hours on a shaker to remove excess dye. OsteoSense (1:100 dilution in 1% BSA/PBS, Revvity, MA, USA) was incubated with the aortic tissue sections for 1 hour at room temperature in the dark. Nuclei were counterstained with DAPI (4’,6’-diamidino-2-phenylindole; 10236276001, Roche, Switzerland) for 5 minutes before mounting (Epredia Shandon Immu-Mount, Thermo Fisher Scientific).

IHC and IF were performed as described [23] (Table 2 and 3). Heat-induced antigen retrieval (HIER) was performed either with citric acid (Vector Laboratories, Inc., USA) or Tris-EDTA pH 9. For IHC, colorimetric development was performed using the ImmPACT® VIP Substrate Kit (Vector Laboratories, CA, USA).

### Myograph measurement

Thoracic aortic pieces (∼4-5 µm in length) were placed in 300 µm pins in the Wire Myograph 620M (DMT, Denmark) for circumferential elasticity measurements. Aortas were stretched until elastin fibers ruptured, indicated by a sudden reduction in holding force. The length change from the initial position until rupture (ΔL) was recorded and compared across treatment groups.

### Electron microscopy & Energy-Dispersive X-ray (EDX) analysis

Transmission electron microscopy (TEM) and EDX analysis were conducted to assess ultrastructural changes and elemental composition in aortic tissues. Sample preparation was performed as described [23]. In short, samples were fixed in 3% glutaraldehyde in 0.1 M Sorensen phosphate buffer, dehydrated in an ethanol series and embedded in epon resin. Sections were examined using a Zeiss Leo 906 transmission electron microscope (Carl Zeiss, Germany).

EDX analyses were performed with the EDAX Genesis system (EDAX, Mahwah, NJ, USA) installed in a FEI XL30 FEG environmental scanning electron microscope (ESEM). The block face was imaged in backscatter mode. Elemental composition (C, O, Na, Cl, P, S, and Ca) was compared between severely and minimally calcified regions within the same tissue section.

### Cell culture & treatment

Mouse aortic smooth muscle cells (MOVAS, ATCC® CRL-2797) were cultured in a growth medium containing DMEM, 10% FBS, 1% Penicillin/Streptomycin, Gentamicin, and Geneticin (G418 Sulfate) (all from Gibco). Cells were maintained under controlled conditions and treated with recombinant mouse PDGF-BB (Sigma-Aldrich, St. Louis, MO) at a concentration of 10 ng/mL for 24 hours to mimic disease conditions. Following treatment, RNA was isolated, and gene expression was analyzed by RT-qPCR as described above.

### Statistical analysis

All statistical analyses were performed using GraphPad Prism 10.4.1 (La Jolla, USA). Data were presented as individual values and mean ± standard deviation (SD) and assessed for normality. Outliers were identified and excluded. For non-normally distributed data, the Mann-Whitney U test was used, while the unpaired two-tailed t-test was applied for normally distributed data comparing two groups. One-way ANOVA with Tukey’s multiple comparisons test was used for comparisons among three or more groups. Two-way ANOVA assessed interactions between two independent variables, followed by Tukey’s test for equal sample sizes or Šídák’s test for unequal sample sizes. Statistical significance was defined as p<0.05.

## Results

### CKD accelerated aortic medial calcification *ex vivo*

We established an *ex vivo* calcification model using murine aortas with known stimulants inducing vascular calcification (Figure 1A) [31]. We first examined the effects of age, sex, genetic background, incubation duration (three to ten days), and aortic region (arch, thoracic, suprarenal, and infrarenal) on calcification. Calcification susceptibility increased with age, peaking between 21 and 30 weeks before reaching a plateau (Supplementary Figure 1A). In contrast, sex and genetic background had no effect (Supplementary Figure 1B-C). Among all incubation periods, the highest degree of Ca^2+^ content was observed at day 10 across all aortic regions (Supplementary Figure 1D-E), with an up to 36-fold increase in aortas cultured in the calcification medium (CM) compared to growth medium (Figure 1B). Aortic medial calcification was confirmed by von Kossa and Alizarin Red staining (Figure 1C). We further validated this finding using transmission electron microscopy (TEM), revealing crystal deposits along the elastin fibers (Figure 1D left). EDX analysis of calcified regions identified a high atomic composition of phosphorus and calcium atoms, corroborating mineralization (Figure 1D right).

**Figure 1.**
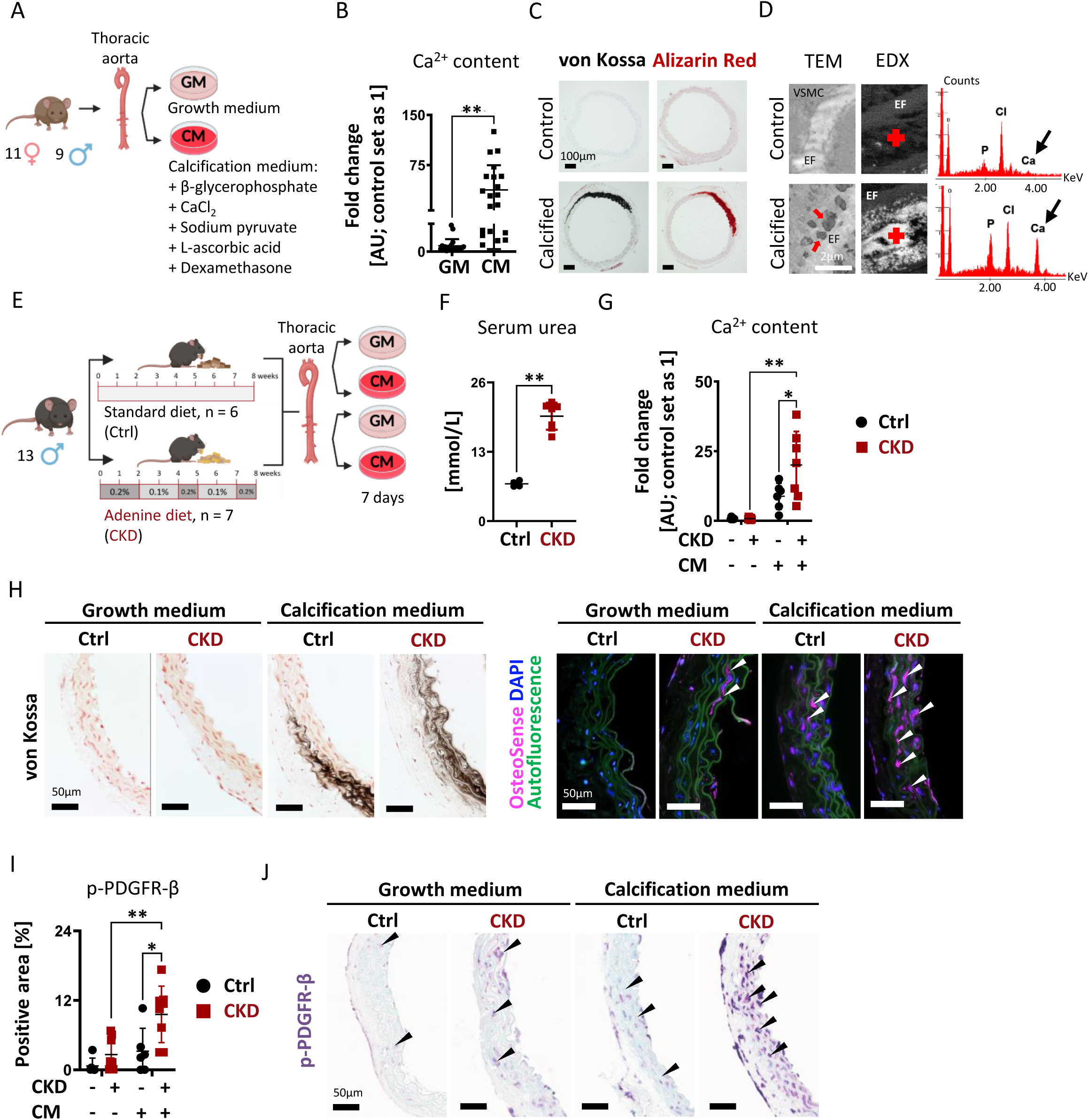
*Ex vivo* aortic calcification was established, showing that aortas from CKD mice (fed an adenine-enriched diet) exhibited increased calcification and PDGFR-β phosphorylation. (A) *Ex vivo* experimental scheme: Aortas were collected from male and female mice and cultured in either growth medium (GM) or calcification medium (CM). CM was supplemented with 10 mmol/L β-glycerophosphate, 8 mmol/L CaCl₂, 10 mmol/L sodium pyruvate, 50 μg/mL L-ascorbic acid, and 100 nmol/L dexamethasone. (B) Calcium content in thoracic aorta tissues was measured, normalized to total protein, and expressed as fold change relative to controls (GM) (set as 1 Arbitrary Unit [AU]). Aortas incubated in CM for 10 days showed significantly increased calcium content. N = 20; **p<0.01. (C) Von Kossa and Alizarin Red S staining highlighted calcified regions as black and red, respectively. Scale bar: 100 µm. (D) Transmission electron microscopy (TEM) images show elastin fibers (EF, white) and VSMCs. The upper image depicts healthy smooth muscle cells, while the lower image shows a calcified aorta with crystal deposits along elastin fibers (red arrows). Scale bar: 2 µm. Energy dispersive X-ray spectroscopy (EDX) analyses confirmed calcification by detecting calcium and phosphorus enrichment (arrow) in highly calcified regions (marked with red plus signs), while chlorine originated from the calcium source (CaCl₂) in the CM. (E) Experimental scheme: Mice were divided into standard diet (Ctrl) and adenine-enriched diet (0.1% and 0.2% adenine) (CKD) groups for 8 weeks. Aortas from these mice were then collected and cultured in GM or CM for 7 days.(F) Serum analysis showed that CKD mice exhibited significantly elevated serum urea levels compared to Ctrl mice, indicating kidney damage. N = 4 for Ctrl, N = 7 for CKD; **p<0.01. (G) After 7 days of *ex vivo* culture, aortas from CKD mice had significantly higher calcium content than those from healthy mice. (H) Von Kossa staining revealed increased calcification in the medial layer of CKD mouse aortas compared to healthy controls. OsteoSense (pink, white arrows) localized to calcified vascular regions, with cell nuclei stained with DAPI (blue) and elastic laminae visible as green autofluorescence. Scale bar: 50 µm. (I) The percentage of area positive for phosphorylated PDGFR-β was significantly higher in calcified CKD mouse aortas compared to healthy controls. N = 6 Ctrl, N = 7 CKD; *p<0.05, **p<0.01. (J) Representative immunohistochemical staining showed greater p-PDGFR-β expression (lilac, black arrows) in aortas from CKD mice compared to control mice. Scale bar: 50 µm. Ctrl: control, GM: growth medium, CM: calcification medium.

Next, we isolated aortas from healthy and CKD mice and subjected them to *ex vivo* calcification conditions (Figure 1E). CKD was confirmed by increased serum urea measurement (Figure 1F). Aortas from CKD mice exhibited significantly increased calcification compared to aortas from healthy animals, with up to 17.8-fold higher Ca^2+^ content (Figure 1G). Visualization of calcification using von Kossa staining revealed a positive signal in the medial region (Figure 1H). OsteoSense staining revealed microcalcifications in the aorta of CKD mice without exposure to calcification medium, which was not detectable by von Kossa staining (Figure 1H). This suggested that CKD alone predisposed aortas to microcalcification.

Immunohistochemistry revealed a significant increase of PDGFR-β phosphorylation in the tunica media of aortas from CKD mice compared to controls when exposed to calcification medium (Figure 1I-J).

### PDGFR-β inhibition attenuated vascular calcification

To determine the role of PDGFR-β in the process of vascular calcification, we treated *ex vivo* calcified aortas with imatinib, a tyrosine kinase receptor inhibitor also targeting PDGFR-β (Figure 2A). Imatinib treatment led to a 2.5-fold reduction in Ca^2+^ content (Figure 2B). Histological analysis confirmed reduced calcification by imatinib treatment (Figure 2C-D).

**Figure 2.**
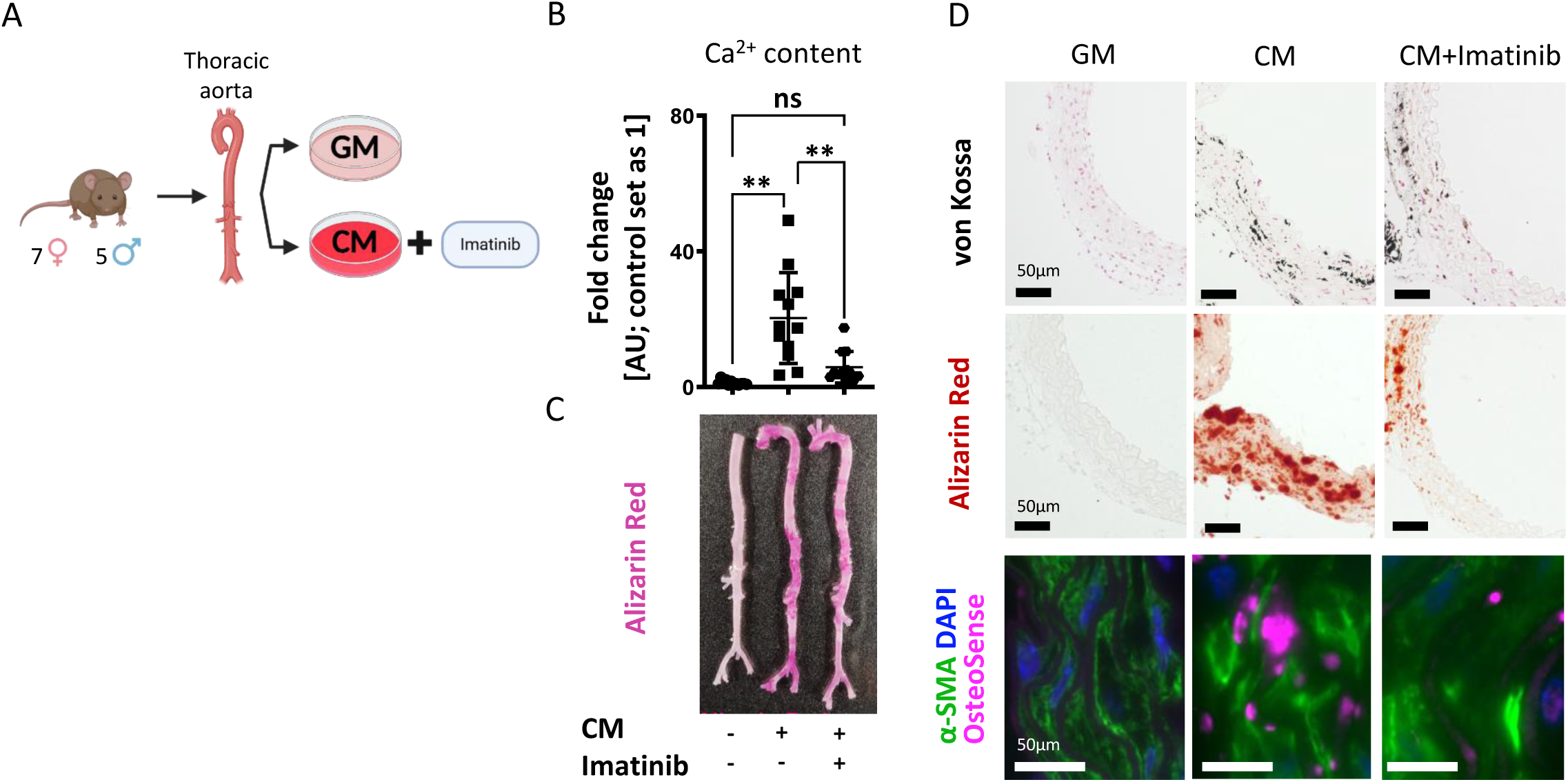
*Ex vivo* imatinib treatment reduces aortic calcification in a non-CKD condition, indicating that PDGFR-β inhibition attenuates calcification. (A) *Ex vivo* experimental scheme: Aortas were harvested from male and female mice and cultured in either growth medium (GM) or calcification medium (CM) supplemented with imatinib (1 µM). (B) Aortas incubated in CM for 10 days resulted in a significant increase in the calcium deposition, which was significantly diminished by the addition of imatinib. N = 12; **p<0.01. (C) Alizarin Red S staining of intact aortas highlighted calcified areas in pink, whereas control aortas remained pale. (D) Histological staining of von Kossa and Alizarin Red S staining revealed calcified regions as black and red, respectively, demonstrating a reduction in calcification upon imatinib treatment. Immunofluorescence staining for α-SMA (green), OsteoSense (pink), and DAPI (blue, nuclei) showed decreased OsteoSense positivity in the imatinib-treated group, further indicating reduced calcification. Scale bar: 50 µm. GM: growth medium, CM: calcification medium.

To model uremic conditions *ex vivo*, we added hemodialysate (HD) to calcification medium (Figure 3A). This resulted in a significant 13.6-fold increase in Ca^2+^ content, highlighting the exacerbating effect of uremia on calcification. Treatment with a soluble PDGFR-β inhibitor, i.e., a recombinant mouse PDGFR-β Fc chimera (sPDGFR-β), led to a significant, 2-fold reduction in Ca^2+^ content in the aorta (Figure 3B; CM vs. CM+HD). Imatinib treatment also significantly attenuated the uremic calcification (Figure 3B).

**Figure 3.**
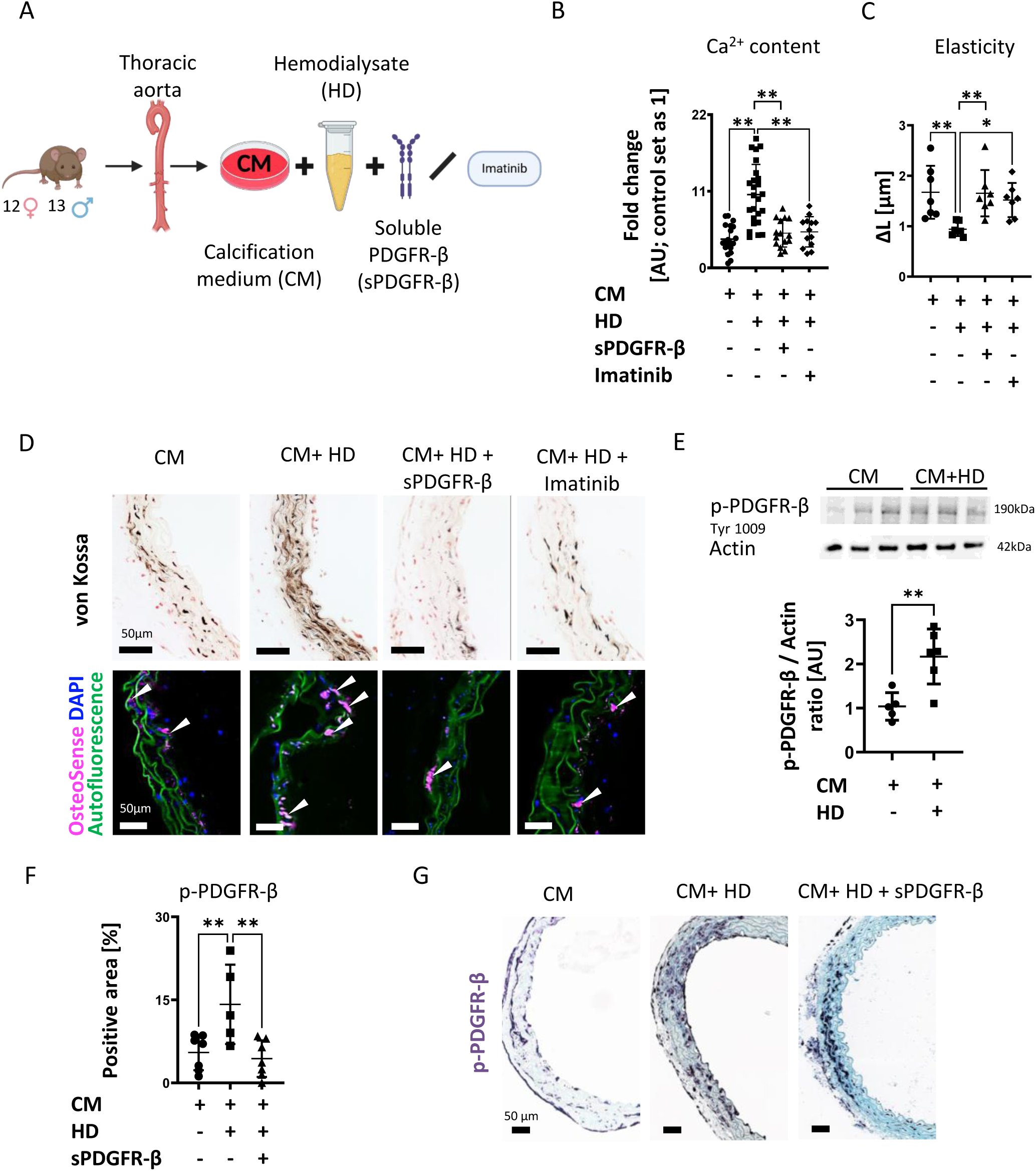
CKD condition, by hemodialysate addition, aggravated aortic calcification, while PDGFR-β inhibition, via soluble PDGFR-β or imatinib, reduced calcification and improved aortic elasticity. (A) Experimental scheme: Hemodialysate (HD) elutes were collected from the wastewater of dialysis patients and added to calcification medium (CM), either alone or in combination with soluble PDGFR-β (0.5 µg/mL) or imatinib (2 µM). (B) Calcium quantification after 7 days of *ex vivo* culture showed that HD addition significantly increased calcification, which was significantly reduced by soluble PDGFR-β or imatinib addition. N = 13–25; **p<0.01. (C) Myograph analysis revealed that the decreased stretch capacity of hemodialysate-treated aortas was restored by soluble PDGFR-β or imatinib addition, improving aortic elasticity. N = 7; **p<0.01. (D) Von Kossa and OsteoSense staining confirmed reduced calcification upon soluble PDGFR-β or imatinib treatment. Cell nuclei were stained with DAPI (blue), and elastic laminae appeared as green autofluorescence. Scale bar: 50 µm. (E) Western blot analysis of p-PDGFR-β (Tyr1009) and actin (loading control) showed increased p-PDGFR-β expression in hemodialysate-treated aortas, indicating that CKD conditions exacerbate PDGFR-β phosphorylation. N = 5–6; **p<0.01.(F) The percentage of area positive for phosphorylated PDGFR-β was significantly reduced after soluble PDGFR-β treatment. N = 5–7; **p<0.01. (F) Representative immunohistochemical staining showed elevated p-PDGFR-β expression in hemodialysate-treated aortas, which was diminished by soluble PDGFR-β addition. Scale bar: 50 µm. CM: calcification medium, HD: hemodialysate, sPDGFR-β: soluble PDGFR-β.

The myograph assay indicated that hemodialysate significantly reduced the aortic elasticity. In line with the Ca^2+^ quantification, both soluble PDGFR-β and imatinib treatment restored elasticity (Figure 3C). Von Kossa and OsteoSense stainings revealed reduced medial calcification in both sPDGFR-β and Imatinib-treated groups (Figure 3D).

At the protein level, Western blots showed that hemodialysate exposure increased PDGFR-β phosphorylation (Figure 3E). This was supported by immunohistochemistry staining, revealing a 1.6-fold increased PDGFR-β phosphorylation, which was significantly reduced by sPDGFR-β treatment (Figure 3F-G).

### Constitutively active PDGFR-β in VSMCs accelerated vascular calcification *ex vivo*

We next generated a genetically modified mouse model with constitutive PDGFR-β activation in the tunica media using the VSMC-specific Myh11 promoter (*Myh11Cre+::Pdgfrb^+/J^*, we term VRbJ). To induce Cre–*loxP* recombination system and activate the mutant *Pdgfrb^+/J^*allele, explanted aortas were treated *ex vivo* with 4-hydroxy tamoxifen. Western blot and immunohistochemistry confirmed nuclear Cre expression (Supplementary Figure 2). Aortas from VRbJ and WT mice were cultured *ex vivo* in growth and calcification medium (Figure 4A). Ca^2+^ content and positive von Kossa arterial media staining were significantly increased in VRbJ compared to WT aortas (Figure 4B-C). The Western blot showed significantly elevated PDGFR-β phosphorylation (2-fold) in VRbJ aortas following calcification medium incubation (Figure 4D-E). These data indicated that enhanced PDGFR-β signaling in VSMCs accelerates calcification. Under uremic conditions, VRbJ aortas also exhibited significantly higher Ca^2+^ content and positive von Kossa staining in the media (Figure 4G-H). Analysis of phosphorylated PDGFR-β confirmed activation of the transgene (Figure 4I-J).

**Figure 4.**
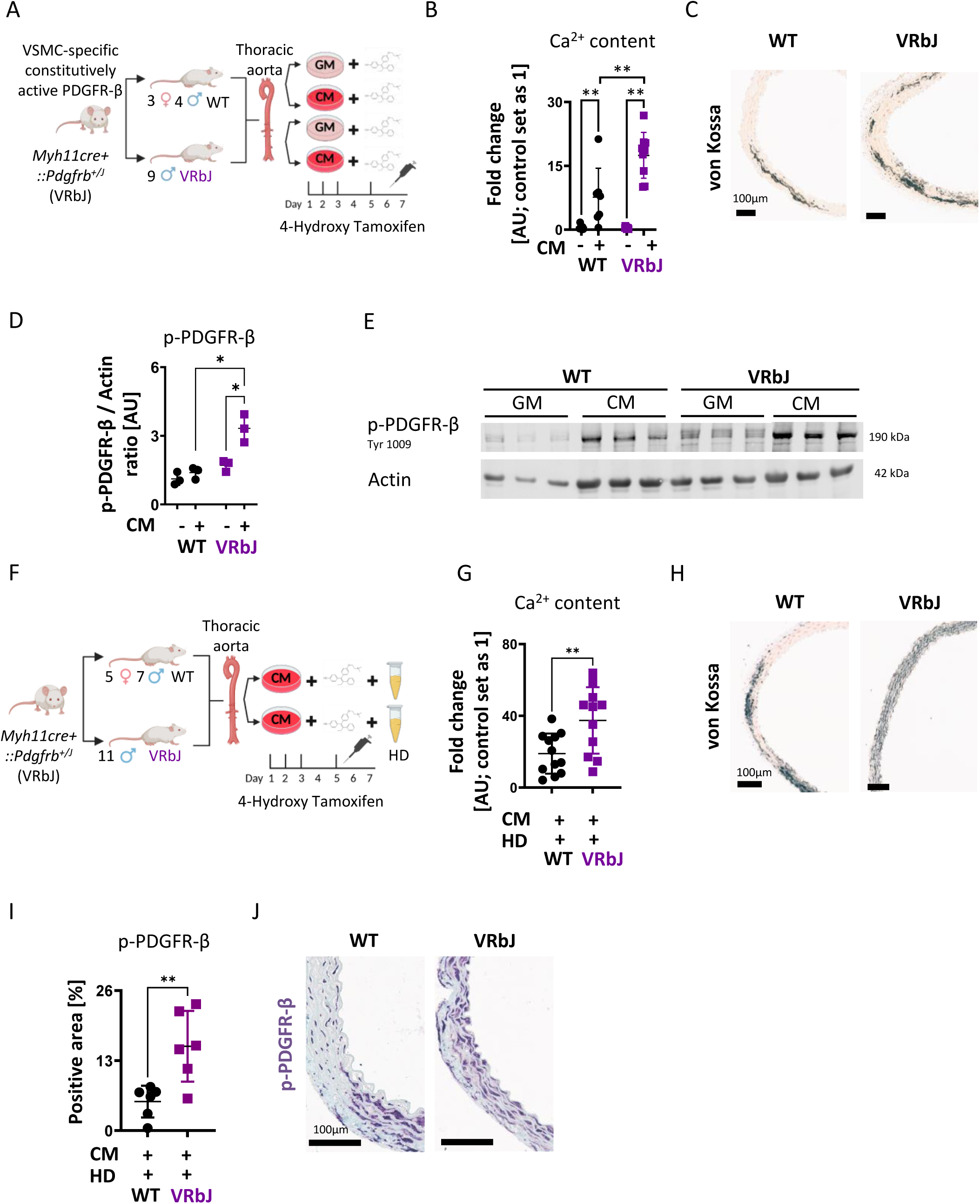
Aortas from constitutively active PDGFR-β (Myh11V536A) mutant mice exhibited increased aortic calcification, particularly under CKD conditions. (A) Experimental scheme: Mice having the PDGFR-β V536A activating mutation in vascular smooth muscle cells (VSMCs) under the Myh11 promoter were generated. Aortas from non-mutant wild type (WT: Myh11Cre-::Pdgfrb^+/+^ and Myh11Cre+::Pdgfrb^+/+^; male and female) and mutant (VRbJ: Myh11Cre+::Pdgfrb^+/J^; male) mice were collected and treated with 4-hydroxy tamoxifen 4 times over 7 days in both growth (GM) and calcification medium (CM). (B) Calcium measurements showed significantly higher calcium content in aortas from VRbJ than WT mice after CM treatment. N = 7 WT, N = 9 VRbJ. (C) Von Kossa staining confirmed increased calcified regions in VRbJ aortas. Scale bar: 100 μm. (D) Densitometric analysis of western blot indicated a slight increase in p-PDGFR-β in control conditions and a significant elevation after CM treatment. N = 3; *p<0.05. (E) Western blot analysis of phosphorylated PDGFR-β (Tyr 1009) normalized to actin in GM and CM-treated groups. N = 3. (F) Experimental scheme: Aortas from WT and VRbJ mice were collected, treated with 4-hydroxy tamoxifen 4 times over 7 days in CM, with the addition of hemodialysate (HD) to mimic CKD conditions. (G) Calcium measurements showed significantly higher content in VRbJ aortas treated with 4-hydroxy tamoxifen, CM, and HD compared to WT controls. (H) Von Kossa staining confirmed more severe calcification in VRbJ aortas, demonstrating that PDGFR-β activation exacerbated calcification under CKD conditions. N = 11–12; **p<0.01. Scale bar: 100 μm. (I) The percentage of area positive for phosphorylated PDGFR-β was significantly higher in VRbJ aortas. N = 6; **p<0.01. (J) Representative histological images showed elevated p-PDGFR-β expression in VRbJ aortas. Scale bar: 100 μm. VSMCs: vascular smooth muscle cells, GM: growth medium, CM: calcification medium, HD: hemodialysate, WT: wild type, VRbJ: V536A mutation in PDGFR-β.

### Constitutively active PDGFR-β in VSMCs accelerated vascular calcification *in vivo*

To investigate the role of PDGFR-β in CKD-associated vascular calcification *in vivo*, we established a mouse model using a PDGFR-β reporter line (*Tg(PDGFRb-eGFP)*). PDGFR-β (GFP) reporter mice were injected with AAV8-D377Y-mPCSK9 viral particles to induce mPCSK9 overexpression and received a mixed diet (Western diet with additional phosphate and adenine) for 12 weeks to induce CKD and associated medial calcification (Supplementary Figure 3A). The aortas of these mice exhibited elevated PDGFR-β as indicated by the positive GFP reporter signal, colocalizing with a positive OsteoSense signal in VSMCs (Supplementary Figure 3B). Quantification of OsteoSense fluorescence revealed a 2.1-fold increase in medial calcification in CKD mice compared to controls (Supplementary Figure 3C). Kidney histology (PAS and AFOG staining, Supplementary Figure 3D) and increased serum creatinine (Supplementary Figure 3G) confirmed the presence of CKD. TEM images revealed structural disruption in the media with elastin fragmentation and deposition of calcium-phosphate crystals (Supplementary Figure 3E). Significant increases in systolic and diastolic blood pressure (Supplementary Figure 3F), inorganic phosphate, and cholesterol levels were also observed (Supplementary Figure 3G). Additionally, hepatic low-density lipoprotein receptor (LDLR) protein levels were significantly decreased (5.2-fold) compared to control mice (Supplementary Figure 3H-I), confirming a successful mPCSK9 overexpression. These data confirmed the successful establishment of uremic medial calcification in PDGFR-β reporter mice.

To dissect the role of PDGFR-β, we induced the CKD calcification model in VRbJ mice and WT littermates (Figure 5A). Both calcifying VRbJ and WT mice did not show significant changes in body weight, systolic and diastolic blood pressure, heart rate, or serum levels of creatinine, cholesterol, inorganic phosphate, and potassium under CKD conditions (Supplementary Figure 4A-E). Creatinine clearance was also comparable between groups after 12 weeks of treatment (Figure 5B), indicating similar CKD severity.

**Figure 5.**
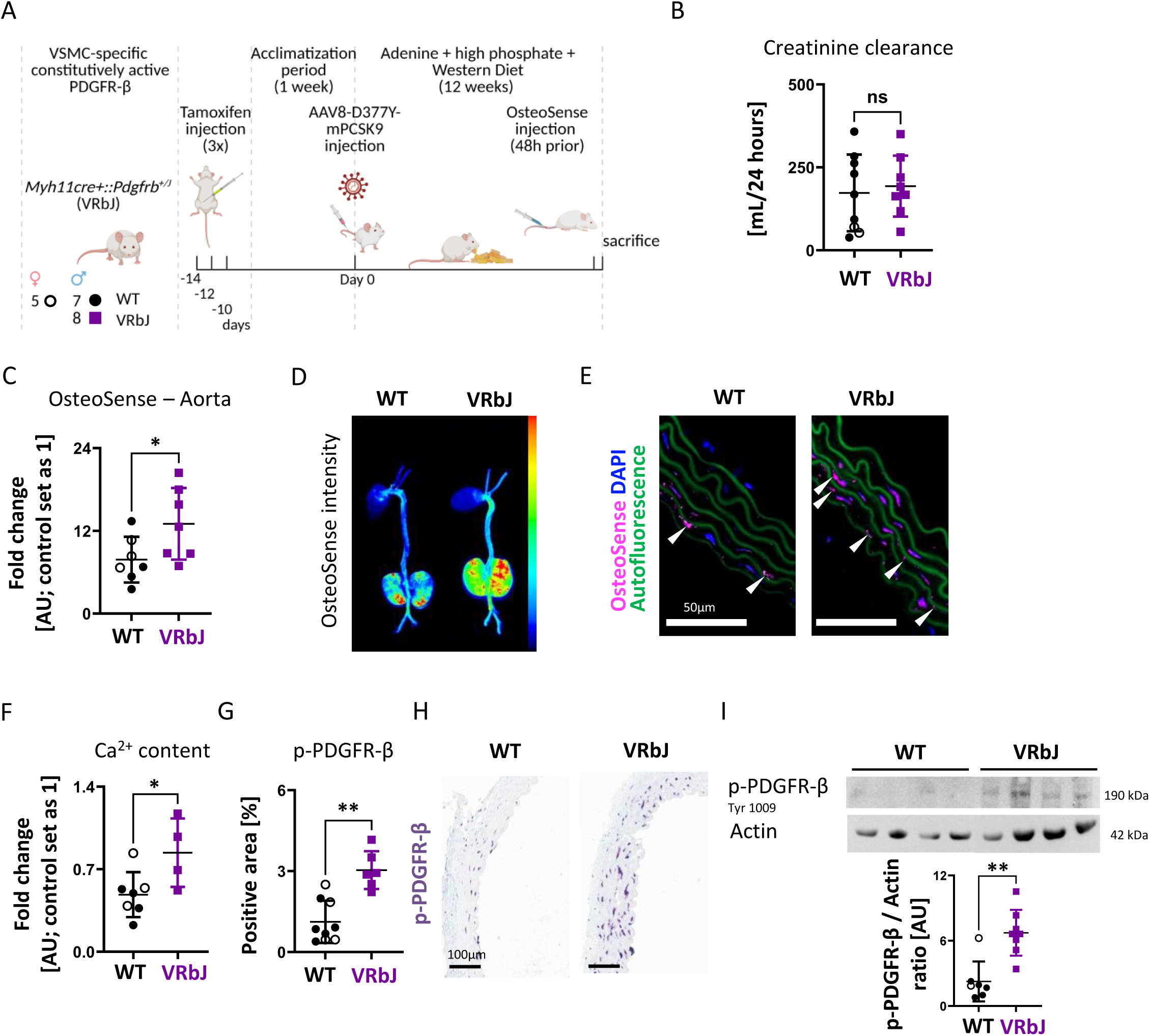
An *in vivo* CKD-vascular calcification model using mPCSK9 injection combined with an adenine, high phosphate (P*i*), and Western diet exacerbated aortic calcification in VSMC-specific constitutively active PDGFR-β mutant mouse model. (A) Experimental design: Mutant (VRbJ: Myh11Cre+::Pdgfrb^+/J^; male) and non-mutant wild-type (WT: Myh11Cre-::Pdgfrb^+/+^ and Myh11Cre+::Pdgfrb^+/+^; male and female) mice received three intraperitoneal tamoxifen injections during the first week (days −14, −12, and −10), followed by a one-week acclimation period. Mice were then injected with AAV8-D377Y-mPCSK9 and maintained on a CKD-inducing diet containing adenine, high phosphate, and a high-fat, high-cholesterol Western diet for 12 weeks. OsteoSense was administered via tail vein injection 48 hours before sacrifice. (B) Serum creatinine clearance measurements indicated no significant difference in kidney function between groups after 12 weeks on the diet. N = 8–9; ns: no significance. (C) Quantification of OsteoSense in aortas, expressed as fold change relative to baseline (set as 1 arbitrary unit [AU]), showed significantly higher calcification in VRbJ aortas. N = 7; *p<0.05. (D) Imaging of the heart, aorta, and kidneys was performed 48 hours after tail vein injection, illustrating increased OsteoSense signal in VRbJ aortas. (E) OsteoSense-positive signals (pink, white arrows) were higher in VRbJ aortas. Nuclei were counterstained with DAPI (blue), and elastic laminae exhibited green autofluorescence. (F) Aortic calcium content was significantly higher in VRbJ aortas compared to WT controls. N = 4–7; *p<0.05. (G) The percentage of area positive for phosphorylated PDGFR-β was significantly higher in VRbJ aortas. N = 6–9; **p<0.01. (H) Representative images of VRbJ aortas demonstrated elevated p-PDGFR-β expression. Scale bar: 100 µm. (I) Western blot analysis of phosphorylated PDGFR-β (Tyr 1009), normalized to actin, showed significantly increased p-PDGFR-β expression in VRbJ aortas compared to WT controls. N = 7–8; **p<0.01. VSMCs: vascular smooth muscle cells, WT: wild type, VRbJ: V536A mutation in PDGFR-β.

OsteoSense was administered 48 hours before sacrifice, allowing *ex vivo* fluorescence imaging of the heart, kidneys and aorta to visualize hydroxyapatite deposition as a marker of calcification (Figure 5D). Compared to WT, aortic calcification was significantly exacerbated in VRbJ mice (Figure 5C), as visualized in thoracic aorta cross-sections (Figure 5E) and further confirmed by Ca^2+^ quantification (Figure 5F). The enhanced aortic calcification was associated with increased phosphorylation of PDGFR-β in VRbJ mice, confirmed by immunohistochemistry and Western blot (Figure 5G-I). Overall, the *in vivo* findings showed aggravated uremic calcification upon PDGFR-β activation in VSMCs.

### PDGFR-β phosphorylation induced a phenotypic switch in VSMC

We treated aortas from VRbJ and WT mice with 4-hydroxy tamoxifen in growth medium for 7 days to investigate downstream effects of PDGFR-β activation, focusing on markers of a calcification-relevant phenotypic switch of VSMCs (Figure 6A). Compared to WT, VRbJ aortas had a significant increase in *Spp1* (osteopontin) and a decrease in *Tagln* (SM22α) expression (Figure 6B), and a 2.4-fold upregulation of osteopontin using immunohistochemistry (Figure 6C-D).

**Figure 6.**
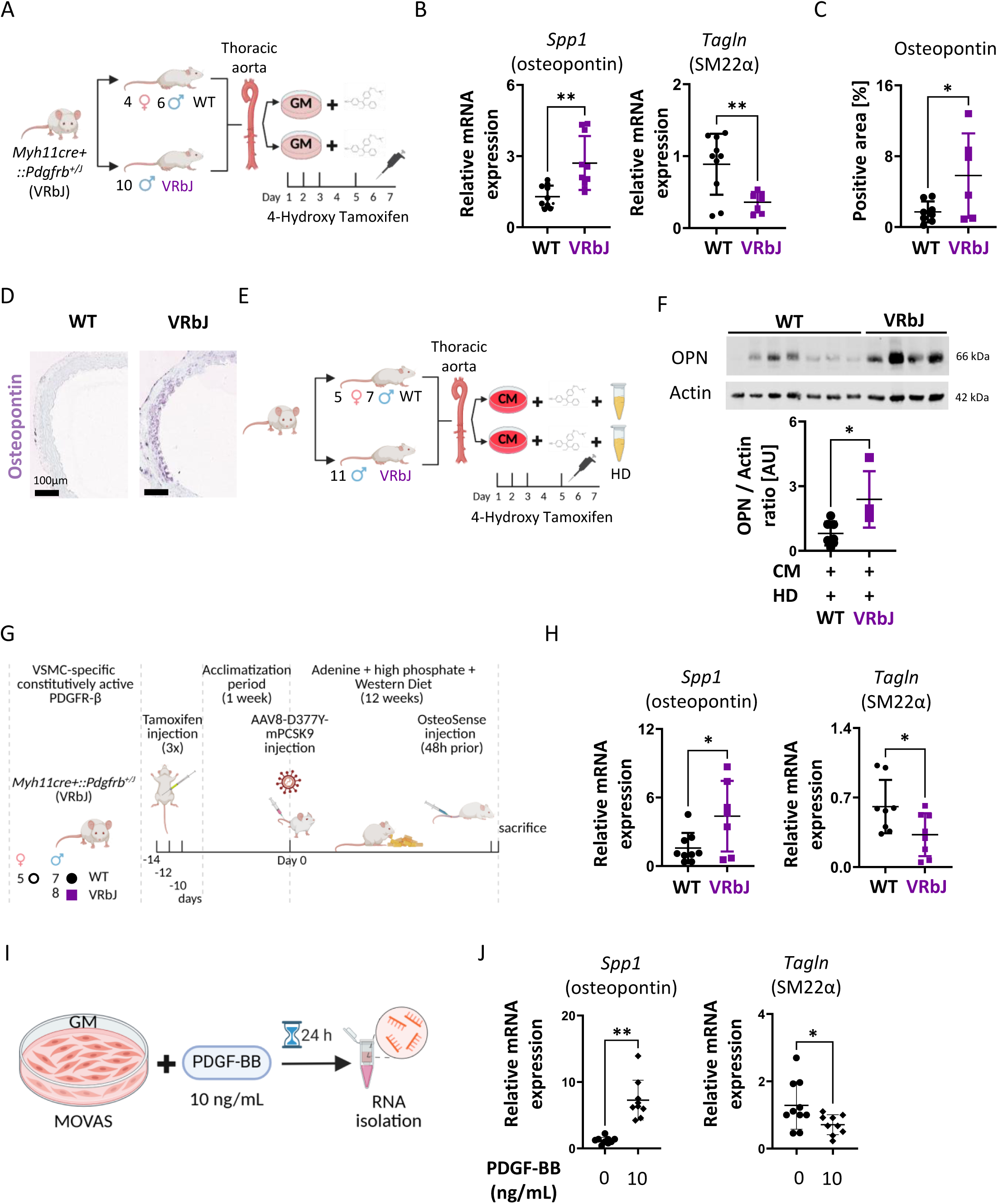
PDGFR-β activation induces a VSMC phenotypic switch by upregulating osteopontin and downregulating SM22α expression. (A) *Ex vivo* experimental scheme: Aortas were collected from non-mutant wild-type (WT: Myh11Cre-::Pdgfrb^+/+^ and Myh11Cre+::Pdgfrb^+/+^; male and female) and mutant (VRbJ: Myh11Cre+::Pdgfrb^+/J^; male) mice and treated with 4-hydroxy tamoxifen four times over seven days in growth medium (GM). (B) Relative mRNA expression of *Spp1* (osteopontin) was significantly increased, whereas *Tagln* (SM22α) expression was significantly reduced in VRbJ aortas compared to WT controls, indicating that PDGFR-β phosphorylation triggered a VSMC phenotypic switch. N = 7–10; **p<0.01. (C) Quantification of VSMCs with elevated osteopontin expression demonstrated a significantly higher percentage in VRbJ aortas. N = 6–8; *p<0.05. (D) Representative immunohistochemical images showed increased osteopontin expression in VRbJ aortas. Scale bar: 100 µm. (E) Experimental scheme: Aortas from WT and VRbJ mice were treated with 4-hydroxy tamoxifen 4 times over 7 days in calcification medium (CM), with hemodialysate (HD) added to simulate CKD conditions. (F) Western blot analysis of osteopontin (OPN), normalized to actin, confirmed significantly higher expression in hemodialysate-treated VRbJ aortas compared to WT aortas. N = 4–7; *p<0.05. (G) *In vivo* experimental scheme: VRbJ and WT mice received three intraperitoneal tamoxifen injections in the first week, followed by one week acclimation period. Mice were then injected with AAV8-D377Y-mPCSK9 and maintained on a CKD-inducing diet for 12 weeks. OsteoSense was administered 48 hours prior to sacrifice via tail vein injection. (H) Relative mRNA levels of *Spp1* were significantly increased, while *Tagln* expression was significantly decreased in VRbJ aortas compared to WT aortas, further confirming the role of PDGFR-β phosphorylation in driving a VSMC phenotypic switch. N = 7–9; *p<0.05. (I) *In vitro* experimental design: MOVAS cells were treated with recombinant PDGF-BB (10 ng/mL) for 24 hours, followed by RNA isolation for gene expression analysis. (J) Quantification of *Spp1* mRNA expression demonstrated a significant upregulation, while *Tagln* showed a significant downregulation upon PDGF-BB treatment, indicating a shift toward osteogenic transdifferentiation. N = 9–10, **p<0.01. CM: calcification medium, HD: hemodialysate, WT: wild type, VRbJ: V536A mutation in PDGFR-β, OPN: osteopontin.

Similar changes were seen in both the *ex vivo* (Figure 6E) and *in vivo* (Figure 6G) uremic calcification models. *Ex vivo*, osteopontin expression was upregulated in VRbJ aorta compared to WT, as shown by Western blot (Figure 6F). *In vivo*, PDGFR-β activation in VRbJ aorta induced a significant upregulation of *Spp1* and downregulation of *Tagln* (Figure 6H).

To further validate these findings, *in vitro* murine smooth muscle cells (MOVAS) were treated with recombinant PDGF-BB (10 ng/mL), a ligand of PDGFR-β, for 24 hours (Figure 6I). This resulted in a significant 5.1-fold increase in *Spp1* expression and a 0.8-fold reduction in *Tagln* expression (Figure 6J), further supporting the role of PDGFR-β in driving osteogenic transdifferentiation.

### PDGFR-β phosphorylation was increased in calcified arteries of CKD patients

Human artery samples with or without calcification were obtained from amputated extremities of patients with and without CKD (Stage 5). The cohort consisted of 21 patients divided into four groups: CKD-calcified (N=5), CKD-non-calcified (N=5), non-CKD-calcified (N=5), and non-CKD-non-calcified (N=6). Patient characteristics are shown in Table 4.

Calcified or non-calcified arteries from non-CKD patients showed comparably low p-PDGFR-β expression in immunohistochemistry (Figure 7A-B). Higher p-PDGFR-β expression was observed in non-calcified arteries from CKD patients, and the highest expression was found in calcified CKD arteries, which was significant compared to calcified non-CKD samples (Figure 7A-B). The p-PDGFR-β expression was not significantly different between calcified and non-calcified arteries within the CKD group, suggesting that CKD *per se* might be sufficient to increase PDGFR-β phosphorylation (Figure 7A-B).

**Figure 7.**
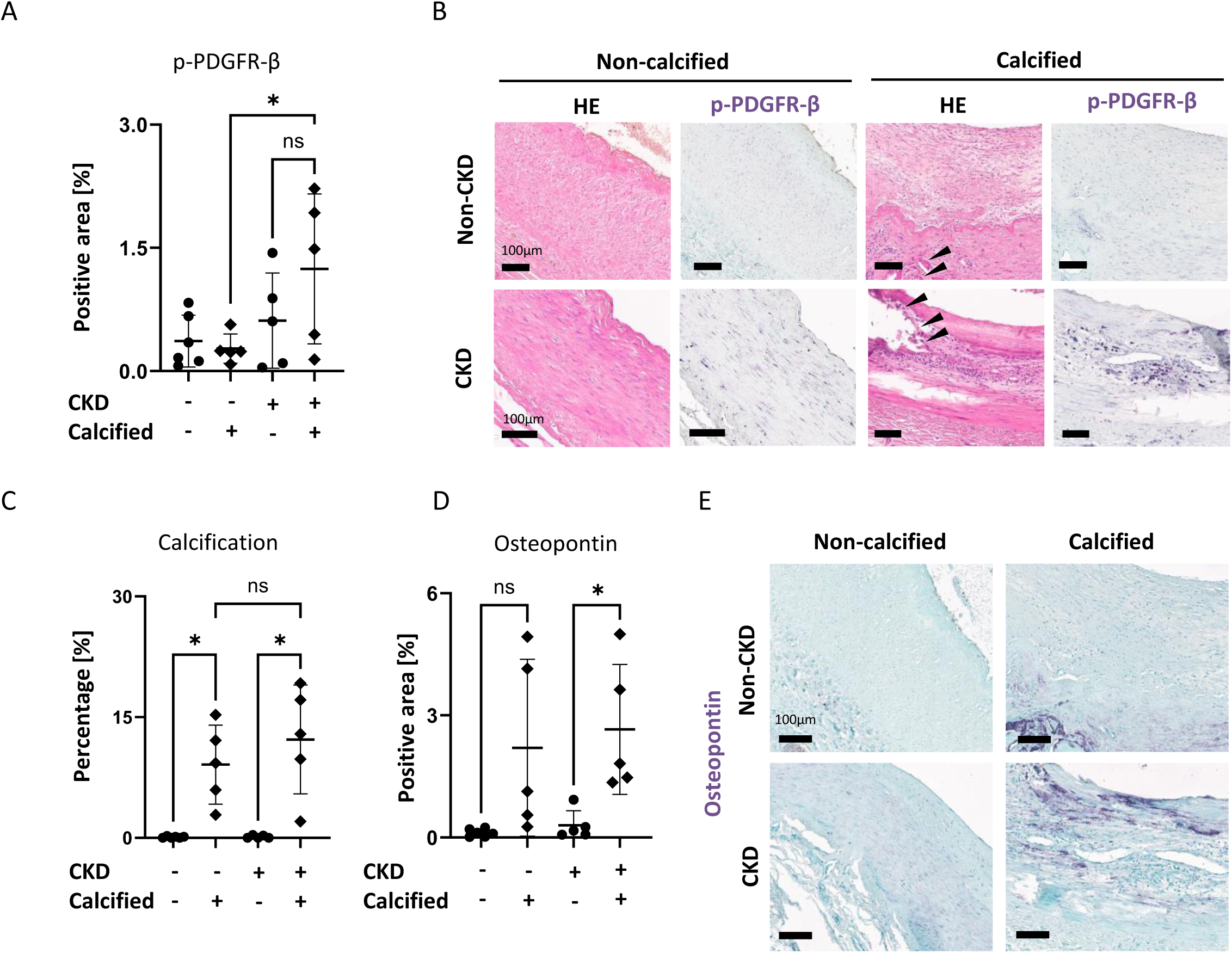
Increased p-PDGFR-β expression observed in calcified arteries compared to non-calcified vessels from CKD and non-CKD patients, together with an upregulation of osteopontin in the calcified arteries. (A) The percentage of area positive for phosphorylated PDGFR-β in the medial layer was significantly higher in calcified aortas from CKD patients compared to calcified vessels from non-CKD patients (N = 5 CKD-calcified, N = 5 CKD-non-calcified, N = 5 non-CKD-calcified, N = 6 non-CKD non-calcified). (B) Representative histological images of hematoxylin and eosin (HE) and phosphorylated PDGFR-β immunostaining in calcified (black arrows) and non-calcified vessels from CKD and non-CKD patients. Scale bar: 100 µm. (C) Quantification of calcified arteries showed no significant difference between CKD and non-CKD. N = 5; ns: no significance. (D) Quantification of osteopontin-positive VSMCs revealed a significantly higher percentage in CKD-calcified arteries compared to CKD-non-calcified arteries. N = 5–6; *p<0.05, ns: no significance. (E) Representative histological images of osteopontin immunostaining in calcified and non-calcified arteries from CKD and non-CKD patients. Scale bar: 100 µm.

Quantitative analysis of histological stainings revealed no significant difference in the percentage of calcified regions between CKD and non-CKD patient arteries in our cohort (Figure 7C).

Osteopontin expression was significantly upregulated (7.9-fold) in calcified arteries compared to non-calcified arteries in CKD patients, with a similar, non-significant trend in non-CKD patients (Figure 7D-E).

## Discussion

Here, we demonstrated that increased PDGFR-β phosphorylation in VSMCs drives the uremic vascular calcification via induction of a phenotypic switch, which could be attenuated by pharmacological and molecular inhibition. To dissect the role of PDGFR-β signaling, we generated a novel transgenic mouse model expressing a constitutively active form of PDGFR-β in VSMCs. Aortas from these mice exhibited enhanced uremic calcification *ex vivo* and *in vivo*, accompanied by upregulation of osteopontin and downregulation of SM22α, i.e. markers of the VSMC phenotypic switch. Human calcified arteries from CKD patients similarly exhibited elevated PDGFR-β phosphorylation and osteopontin content. To our knowledge, this is the first study to directly link PDGFR-β activation to osteopontin expression in the context of vascular calcification. Collectively, our findings identify PDGFR-β as a mechanistic driver and potential therapeutic target in CKD-associated vascular pathology.

The PDGF system is well studied in VSMCs, particularly in the context of vascular development [32]. Beyond its developmental role, PDGFR-β activation has been implicated in vascular pathology through inducing inflammation, oxidative stress, and phenotypic modulation of VSMCs [17]. Activated PDGFR-β has been shown to induce cerebral microvascular calcification [33]. Enhanced PDGFR-β signaling driven by an activating mutation also amplifies and accelerates atherosclerosis by promoting advanced plaque development and ectopic lesion formation in the thoracic aorta and coronary arteries [34]. Although VSMC-specific activation of PDGFR-β via an SM22α-Cre system induced dedifferentiation and medial thickening [18], that study did not evaluate microcalcification. Our findings extend this work by providing direct evidence that PDGFR-β activation contributes to vascular calcification via phenotypic switching of VSMCs.

PDGF-BB, one activator for PDGFR-β, is a well-established driver of VSMC migration, proliferation, extracellular matrix production, and phenotypic switching, primarily through the downregulation of contractile markers and induction of a synthetic phenotype [35]. During this transition, VSMCs lose lineage-specific markers such as SM22α and acquire osteogenic features, including upregulation of osteopontin, thereby contributing to vascular calcification [36]. Notably, PDGF-BB can also bind to PDGFR-α, making it unclear in ligand-based studies which receptor primarily mediates the phenotypic switch. While PDGFR-β has been implicated in promoting this phenotypic transformation [37], the underlying mechanisms remain incompletely defined. Using our tamoxifen-inducible transgenic mouse model with constitutive PDGFR-β activation in VSMCs, we revealed that PDGFR-β activation alone was sufficient to induce a VSMC phenotypic switch in the absence of calcification medium, underscoring its upstream role in osteogenic signaling.

Our cohort of non-calcified and calcified arteries from non-CKD and CKD patients showed an upregulating trend of PDGFR-β phosphorylation in CKD arteries, regardless of calcification status, suggesting that CKD alone may activate PDGFR-β signaling. Consistently, in our *ex vivo* model, aortas from CKD mice cultured in growth medium exhibited early-stage microcalcifications detectable by OsteoSense but not by von Kossa staining. These vessels also showed PDGFR-β phosphorylation levels comparable to calcified non-CKD aortas, indicating that the uremic milieu can prime the vasculature for calcification via PDGFR-β activation, even in the absence of overt calcific stimuli.

In both *ex vivo* and *in vivo* models, PDGFR-β activation was associated with decreased expression of *Tagln* (SM22α) and increased expression of *Spp1* (osteopontin), markers of VSMC phenotypic switching. Notably, in our human cohort, phosphorylated PDGFR-β and osteopontin were most prominently expressed in calcified arteries of CKD patients, reinforcing a mechanistic link between PDGFR-β activation and osteopontin upregulation in CKD. These findings suggest that PDGFR-β activation may precede and potentially drive the osteogenic reprogramming of VSMCs in CKD, even in the absence of overt calcification. We propose that the CKD milieu may act as a priming factor by inducing PDGFR-β phosphorylation, thereby promoting a switch to an osteogenic phenotype and increasing susceptibility to vascular calcification. To our knowledge, this is the first study to directly demonstrate a causal link between PDGFR-β activation and VSMC phenotypic switching, independent of a calcific stimulus.

PDGFR-β phosphorylation can be effectively inhibited by imatinib, a tyrosine kinase inhibitor, which has shown efficacy in reducing venous stenosis in vascular grafts [38], *in vivo* neointimal formation following vascular injury [39], and repressing vascular remodeling in pulmonary hypertension [40]. Other PDGFR-β–targeted strategies, including AG-1295 and the RNA aptamer Apt 14, have similarly attenuated neointimal hyperplasia in preclinical models [41, 42]. More recently, the randomized, double-blind Phase 2 TORREY trial evaluated seralutinib, an inhaled kinase inhibitor targeting PDGFR-β (alongside CSF1R and c-KIT), in adults with pulmonary arterial hypertension and showed a significant reduction in pulmonary vascular resistance [43]. While these studies primarily focused on mitigating VSMC and endothelial cell proliferation, migration, and vascular remodeling, their relevance to vascular calcification remained unexplored. Here, we show that PDGFR-β inhibition via imatinib or soluble PDGFR-β robustly reduced calcification under both uremic and non-uremic conditions in an *ex vivo* model. Notably, PDGFR-β inhibition, via either soluble PDGFR-β or imatinib, has also been shown to mitigate kidney fibrosis in experimental models of unilateral ureteral obstruction and ischemia–reperfusion injury [44], suggesting that systemic PDGFR-β inhibition may have dual benefit by both attenuating CKD progression and limiting vascular calcification.

Multi-omics approaches integrating genomics, transcriptomics, proteomics, and metabolomics have identified *SPP1* (osteopontin) as a key regulator of vascular calcification, with a central position in protein–protein interaction networks [45]. Proteomic profiling using a multiplex biomarker panel in CKD further confirmed osteopontin as an early marker of CKD progression [46]. Notably, osteopontin levels were associated with inflammatory cytokines and bone metabolism markers, reflecting the severity of vascular pathology and predicting disease advancement [46]. These data align with our findings that PDGFR-β–driven osteopontin upregulation may represent a mechanistic link between CKD and vascular calcification.

While our findings are supported by complementary *ex vivo*, *in vivo*, and human data, the causal role of PDGFR-β in human vascular calcification remains inferential. PDGFR-β inhibitors, Imatinib and soluble PDGFR-β, may exert off-target effects or impact other signaling pathways, which might limit the specificity of PDGFR-β–directed interventions. All mutant mice were male, precluding assessment of sex-specific responses. Additionally, although our human cohort included both CKD and non-CKD arteries, the limited sample size and heterogeneity in vascular bed origin may impact the generalizability of our findings. Importantly, only patients with stage 5 CKD were included, which limits our ability to determine whether PDGFR-β phosphorylation is an early event in CKD progression that precedes vascular calcification. At this advanced stage, medial calcification was already present in both CKD and non-CKD arteries. Future studies involving earlier CKD stages, larger, sex-balanced cohorts, and more selective PDGFR-β inhibitors will be essential to validate and extend the translational relevance of our findings.

In conclusion, our study identified PDGFR-β as a key mediator of VSMC phenotypic switching and vascular calcification, particularly under CKD conditions. Through a combination of *ex vivo*, *in vivo*, and human *ex vivo* analyses, we demonstrated that PDGFR-β activation promoted osteogenic reprogramming of VSMCs even in the absence of calcific stimuli, while its inhibition markedly attenuated calcification. These findings provide new mechanistic insights and identify PDGFR-β as a promising therapeutic target for vascular calcification in both CKD and non-CKD populations. Given the limited treatment options for vascular calcification in CKD, our results support further investigation into PDGFR-β inhibition as a novel intervention strategy.

## Acknowledgments

We thank Louisa Böttcher, Pinar Sönmez, Christina Gianussis, Jana Baues, and Marie Cherelle Timm for excellent technical assistance, and Julia Peusquens and Alexander Slowik for support with animal documentation.

The study was supported by the German Research Foundation (DFG, Project IDs 322900939 & 445703531), European Research Council (ERC Consolidator Grant No 101001791), and the Federal Ministry of Education and Research (BMBF, STOP-FSGS-01GM2202C), all to P.B.

## Author Contributions

B.Y.Ö., D.W.L.W., B.M.K., and P.B. conceived and supervised the study. B.Y.Ö., D.W.L.W., and L.Z. performed the experiments. D.W.L.W. oversaw animal breeding and regulatory protocols and co-authored the animal application with B.M.K., B.Y.Ö. and D.W.L.W. conducted *in vivo* animal experiments. S.v.S. provided the human artery cohort; L.O. contributed PDGFR-β mutant mice; J.M. provided CKD animals for *ex vivo* studies. J.J. and V.J. supplied hemodialysate; E.M.B. performed TEM analysis. H.N. and C.G. provided materials and technical support. P.D., M.H., N.M., H.N. and J.F. contributed conceptual input. B.Y.Ö. conducted statistical analyses, prepared figures, and wrote the first draft of the manuscript. D.W.L.W., S.v.S., B.M.K., and P.B. critically revised the manuscript. All authors reviewed and approved the final version. B.Y.Ö., D.W.L.W., and P.B. are guarantors of the study and take full responsibility for the integrity of the data and the accuracy of the analysis.

## Supplementary Figures

**Supplementary Figure 1.**
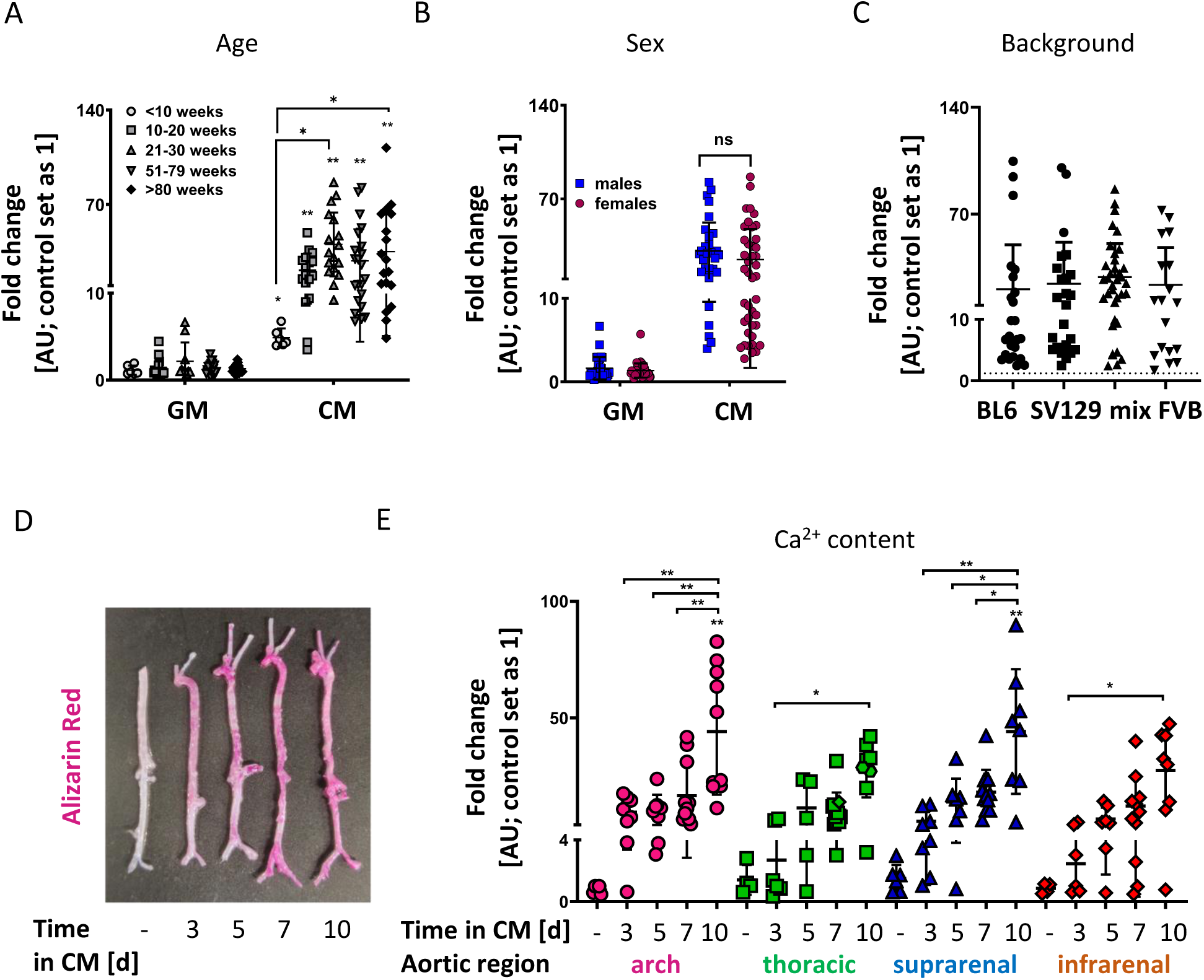
Effect of incubation time, aortic region, age, sex, and mouse genetic background on *ex vivo* vascular calcification. (A) Calcium measurements in mice of different ages demonstrated that calcification increased with age up to 30 weeks, with no further progression beyond this point. All age groups exhibited a significantly higher calcium content compared to their control group, as indicated above the respective values. N = 6–22. *p<0.05, **p<0.01. (B) No significant difference in calcification levels was observed between male (N = 32) and female (N = 42) mice. ns: no significance. (C) Calcium measurements across different genetic backgrounds (C57BL/6J, SV129, a mixed C57BL/6-SV129 strain, and FVB) showed no significant differences in calcification. The dashed line represents control group values. N = 18–36. (D) Alizarin Red S staining of whole aortas after incubation in growth medium (control, 10 days) or calcification medium (CM) for 3, 5, 7, or 10 days. Pink areas indicate calcified regions. (E) Calcium measurements from different aortic regions (arch, thoracic, suprarenal, and infrarenal) at various incubation times (3-10 days) showed that prolonged exposure to calcification medium resulted in increased calcium content across all regions. N = 4–12; *p<0.05, **p<0.01. GM: growth medium, CM: calcification medium.

**Supplementary Figure 2.**
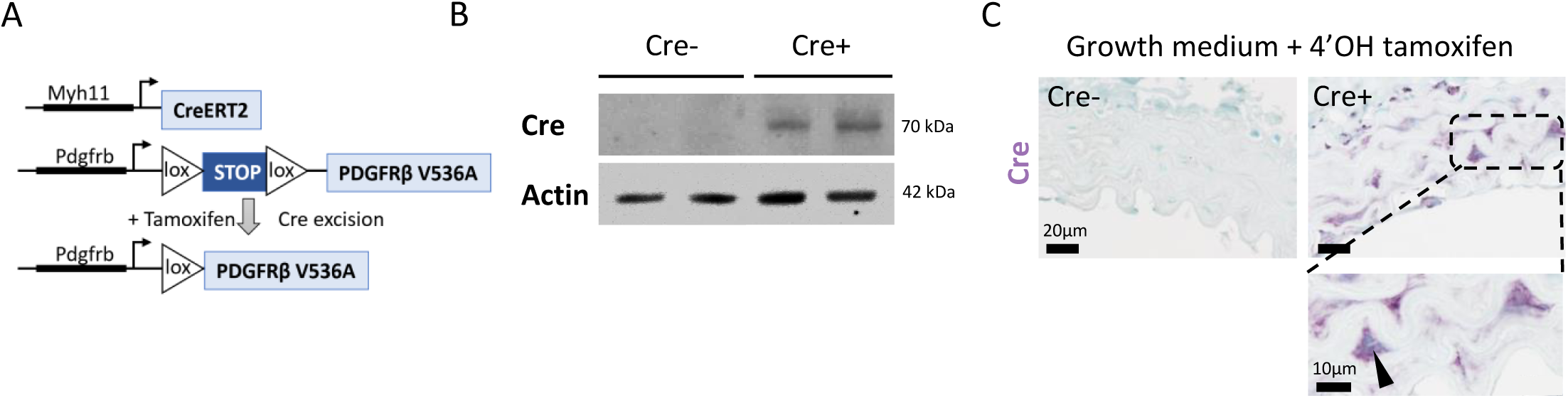
E*x vivo* analysis of aortas from Myh11creER^T2^ mice treated with 4-hydroxy tamoxifen. (A) Genetic strategy: Schematic representation of the conditional activation of the PDGFR-β V536A allele in vascular smooth muscle cells (VSMCs) under the control of the *Myh11* promoter. Tamoxifen administration activates Cre recombinase, excising the loxP-flanked stop cassette and allowing transcription of full-length *Pdgfrb* cDNA containing a single valine-to-alanine substitution at residue 536 (V536A), introduced via a knock-in strategy. (B) Western blot analysis confirmed Cre recombinase protein expression in Cre+ aortas after 4-hydroxy tamoxifen treatment, while absent in Cre− aortas. Actin served as a loading control. (C) Representative immunohistochemical staining showed Cre expression (purple) in the medial layer of Cre+ aortas, with nuclear localization (arrow) confirming successful activation of the Cre-lox system. Scale bars: 20 µm and 10 µm.

**Supplementary Figure 3.**
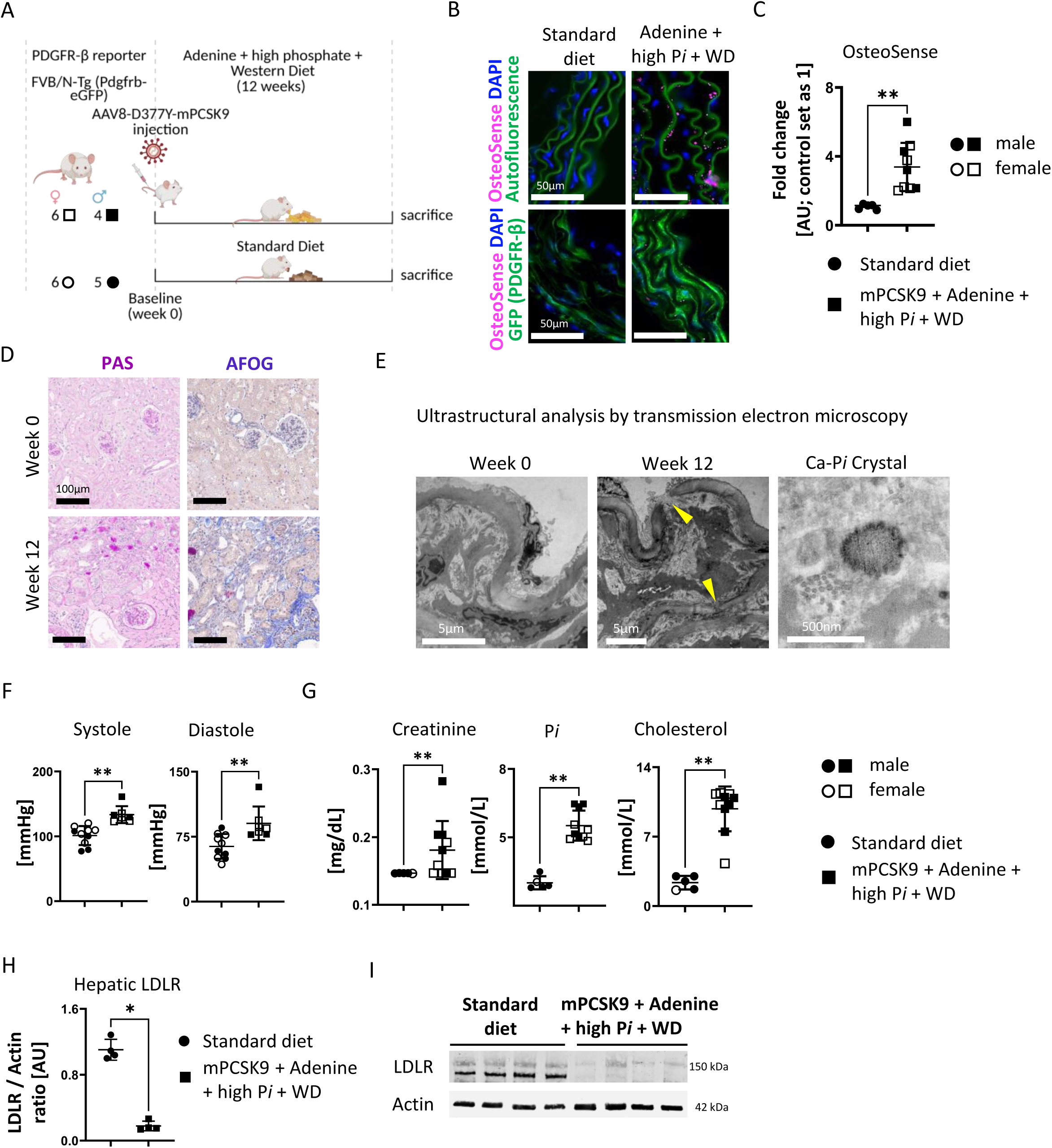
*In vivo* CKD-vascular calcification model using mPCSK9 injection combined with adenine, high phosphate (P*i*), and Western diet induced PDGFR-β expression and exacerbated calcification. (A) Schematic of the *in vivo* experimental model: Male and female PDGFR-β reporter mice (FVB/N-Tg Pdgfrb-eGFP) were injected with AAV8-D377Y-mPCSK9 to induce PCSK9 overexpression and maintained on a mixed diet containing adenine, high phosphate, and a Western diet for 12 weeks. (B) OsteoSense-positive signals (pink) were observed exclusively in CKD aortas, confirming vascular calcification. Nuclei were counterstained with DAPI (blue), and elastic laminae exhibited green autofluorescence (upper images). Lower images demonstrated increased PDGFR-β expression (GFP) in aortas from CKD mice. Scale bar: 50 µm. (C) Quantification of OsteoSense intensity, expressed as fold change relative to controls (standard diet-fed mice, set as 1 Arbitrary Unit [AU]), demonstrated significantly higher OsteoSense positivity in CKD aortas. N = 5 control, N = 9 CKD; **p<0.01. (D) Representative histological images (PAS: Periodic Acid–Schiff and AFOG: Acid Fuchsin Orange G) revealed kidney damage in CKD mice. Scale bar: 100 µm. (E) Ultrastructural analysis of the aortic medial layer by transmission electron microscopy revealed structural disruption in CKD mice, with discontinuous elastin fibers (yellow arrows) and a precipitated calcium phosphate crystal. Scale bars: 5 µm, 500 nm. (F) CODA tail-cuff measurements demonstrated significantly elevated systolic and diastolic blood pressure in CKD mice post-treatment. N = 7-10. **p<0.01. (G) Serum analysis showed a significant increase in creatinine, inorganic phosphate, and cholesterol levels in CKD mice compared to baseline. N = 5-10. **p<0.01. (H) Densitometric analysis of western blot demonstrated a significant reduction in hepatic low-density lipoprotein receptor (LDLR) expression following viral injection, confirming mPCSK9 overexpression. N = 4; *p<0.05. (I) Western blot analysis of LDLR, normalized to actin, was performed in standard diet and mixed diet with mPCSK9-treated groups. N = 4 per group.

**Supplementary Figure 4.**
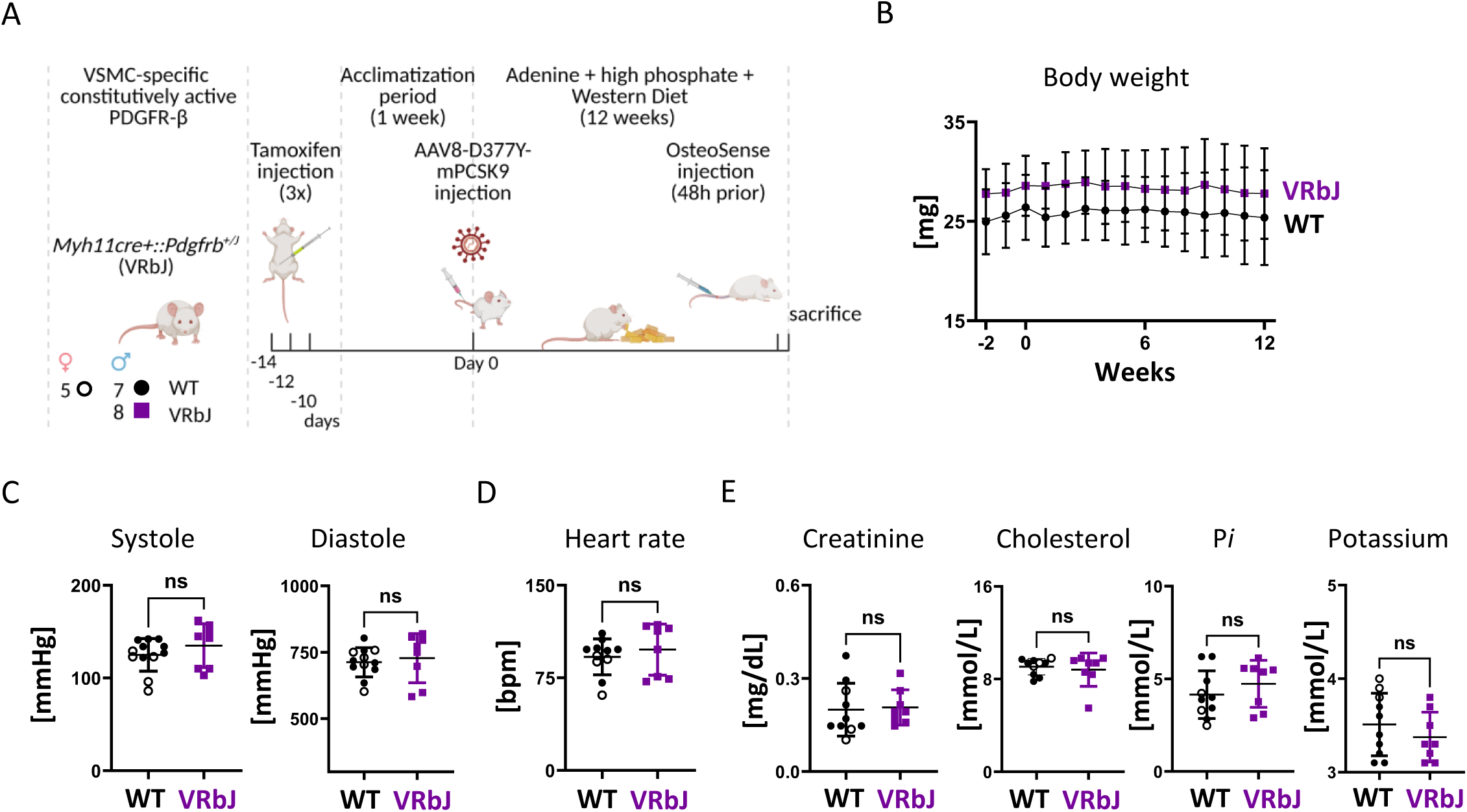
*In vivo* analysis of PDGFR-β mutant (VRbJ) mice revealed similar levels of kidney and cardiovascular dysfunction following 12 weeks of a mixed diet (adenine + high Pi + Western diet) compared to non-mutant (WT) mice. (A) *In vivo* experimental scheme: Mutant (VRbJ: Myh11Cre+::Pdgfrb^+/J^; male) and non-mutant wild-type (WT: Myh11Cre-::Pdgfrb^+/+^ and Myh11Cre+::Pdgfrb^+/+^; male and female) mice received three intraperitoneal tamoxifen injections in the first week (days −14, −12, and −10), followed by a one-week acclimation period. Mice were then injected with AAV8-D377Y-mPCSK9 and maintained on a mixed diet for 12 weeks. 48 hours prior to sacrifice, they were injected OsteoSense via tail vein. (B) Body weight did not significantly differ between VRbJ and WT mice throughout the study. N = 8-12. (C) Systolic and diastolic blood pressure measurements showed no significant differences between VRbJ and WT mice. N = 8-12. ns: no significance. (D) Heart rate remained comparable between groups. N = 8-11. ns: no significance. (E) Serum analysis showed no significant difference in creatinine, cholesterol inorganic phosphate, or potassium levels between VRbJ and WT mice, indicating comparable levels of kidney and cardiovascular dysfunction. N = 8-10. ns: no significance.

